# Large-scale conformational changes of FhaC provide insights into the two-partner secretion mechanism

**DOI:** 10.1101/2021.11.09.467682

**Authors:** G. Sicoli, A. Konijnenberg, J. Guerin, S. Hessmann, E. Del Nero, O. Hernandez-Alba, S. Lecher, G. Rouaut, L. Müggenburg, H. Vezin, S. Cianférani, F. Sobott, R. Schneider, F. Jacob-Dubuisson

## Abstract

The Two-Partner secretion pathway mediates protein transport across the outer membrane of Gram-negative bacteria. TpsB transporters belong to the Omp85 superfamily, whose members catalyze protein insertion into, or translocation across membranes without external energy sources. They are composed of a transmembrane β barrel preceded by two periplasmic POTRA domains that bind the incoming protein substrate. Here we used an integrative approach combining *in vivo* assays, mass spectrometry, nuclear magnetic resonance and electron paramagnetic resonance techniques suitable to detect minor states in heterogeneous populations, to explore transient conformers of the TpsB transporter FhaC. This revealed substantial, spontaneous conformational changes with a portion of the POTRA2 domain coming close to the lipid bilayer and surface loops. Specifically, the amphipathic β hairpin immediately preceding the first barrel strand can insert into the β barrel. We propose that these motions enlarge the channel and hoist the substrate into it for secretion. An anchor region at the interface of the β barrel and the POTRA2 domain stabilizes the transporter in the course of secretion. Our data propose a solution to the conundrum how these transporters mediate protein secretion without the need for cofactors, by utilizing intrinsic protein dynamics.

## Introduction

The Two-Partner Secretion (TPS) pathway is dedicated to the export of large proteins notably serving as virulence factors (Guerin et al., 2017). The TpsB transporters are transmembrane β-barrel proteins that secrete their substrates, collectively called TpsA proteins, across the outer membrane of various Gram-negative bacteria. They belong to the ubiquitous Omp85 superfamily whose members mediate protein insertion into, or translocation across membranes of bacteria and eukaryotic organelles, and which includes the essential bacterial BamA transporters (Heinz & Lithgow, 2014; Knowles et al., 2009; Noinaj et al., 2017). The FhaB/FhaC pair of *Bordetella pertussis* is a model TPS system, in which the FhaC transporter mediates the translocation of the adhesin FhaB across the outer membrane (Fan et al., 2012).

Omp85 transporters are composed of N-terminal POTRA (<u>po</u>lypeptide <u>tr</u>ansport <u>a</u>ssociated) domains - two in the case of TpsB transporters - followed by a 16-stranded transmembrane β barrel, which for FhaC is the FhaB translocation pore (Baud et al., 2014). The POTRA domains mediate protein-protein interactions in the periplasm, and notably recognition of client proteins (Delattre et al., 2011). Another hallmark feature of the Omp85 superfamily is the extracellular loop L6 that folds back inside the barrel and harbors a conserved motif at its tip forming a salt bridge interaction with a specific motif of the inner β-barrel wall (Gu et al., 2016; Maier et al., 2015; Noinaj et al., 2013).

A specific feature of TpsB transporters is an N-terminal α helix called H1 that plugs the β barrel (Clantin et al., 2007; Guerin et al., 2014; Guerin et al., 2020; Maier et al., 2015) (Fig. 1A, B). An extended linker follows H1 and joins it to the POTRA1 domain in the periplasm. Recently, the X-ray structures of the TpsB transporters CdiB^Ab^ and CdiB^Ec^ have shown very similar folds to that of FhaC, albeit with slightly different positions of H1 in the barrel (Guerin et al., 2020). Both H1 and L6 stabilize the barrel in a closed conformation that most likely corresponds to the resting state of the transporter (Guerin et al., 2020; Maier et al., 2015). The β barrel, the L6 loop and the two POTRA domains are essential for transport activity (Clantin et al., 2007).

**Figure 1.**
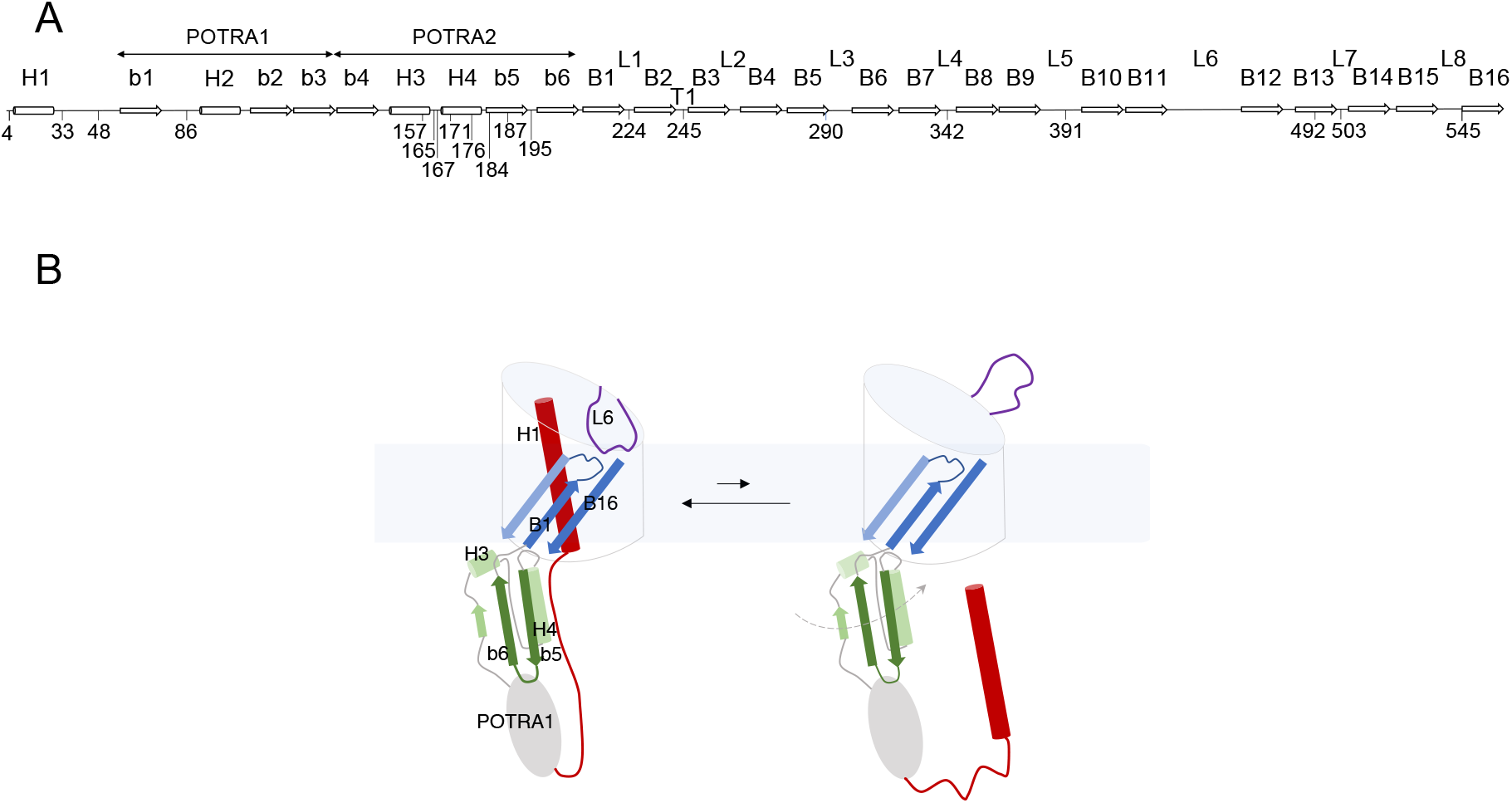
FhaC: Secondary structure and intrinsic dynamics. **(A)** Linear representation of the secondary structure elements of FhaC, with residues used in this work. L1 to L8 represent the extracellular loops, b1 to b6 the *β* strands of the POTRA domains, H1 to H4 the *α* helices, and B1 to B16 the *β*-barrel strands. T1 is the first periplasmic turn. (B) Schematic representation of known FhaC motions linked to its activity. Specific structural elements are colored as follows: H1 and the linker in red, the POTRA2 domain in shades of green, L6 in purple, and *β* barrel strands B1, B2 and B16 in shades of blue. In its resting conformation (left), FhaC is closed, with H1 crossing the *β*barrel. The linker is positioned along the H4 helix of the POTRA2 domain (Maier et al., 2015), thus obstructing a major substrate binding site (Delattre et al., 2011). L6 is folded inside the barrel with a conserved interaction between its tip and the barrel wall. This conformation is in equilibrium with an open form (right), in which H1 has vacated the pore and the substrate binding site is available (Guerin et al., 2014). We have obtained evidence that L6 also moves away from its resting position in the *β* barrel and that the POTRA2 domain undergoes so far undefined motions (represented as a dashed line) (Guerin et al., 2015).

Omp85 transporters likely function in the absence of ATP or an electrochemical gradient. They appear to be very dynamic and to undergo conformational cycling (Doerner & Sousa, 2017; Guerin et al., 2020; Hartmann et al., 2018; Iadanza et al., 2020; Renault et al., 2011; Warner et al., 2017). Lateral opening of the barrel between the first and last anti-parallel β strands is a common mechanistic feature of Omp85 transporters, which is involved in their respective functions (Diederichs et al., 2020; Doyle & Bernstein, 2019; Estrada Mallarino et al., 2015; Guerin et al., 2020; Höhr et al., 2018; Iadanza et al., 2016; Noinaj et al., 2014; Tomasek et al., 2020).

The mechanism of two-partner secretion remains poorly understood, but it is known to involve substantial conformational changes of the transporter including exit of H1 from the β barrel and motions of the L6 loop (Guerin et al., 2014; Guerin et al., 2020; Guerin et al., 2015) (Figure 1B). The motion of H1 toward the periplasm is facilitated by conformational changes of flexible regions of the barrel, in particular the first β-barrel strand B1 and the extracellular loops L1, L2 and L6 (Guerin et al., 2020). Binding of the N-terminal, conserved ‘TPS’ domain of the substrate protein to the POTRA domains of its transporter also appears to enhance conformational changes (Guerin et al., 2015). How the substrate enters the pore and is progressively hoisted towards the cell surface without backsliding to the periplasm remains unknown, but we hypothesize a mechanism implying yet uncharacterized transient conformations of TpsB transporters. In this work we have explored such FhaC conformers using biophysical techniques suitable to detect minor states in heterogeneous populations. Our data revealed an intrinsic exchange of the POTRA2 domain between several conformations in slow equilibrium, and that these conformational changes are linked to transport activity.

## Results

### Effects of freezing the POTRA2 conformation on secretion activity

We have previously obtained evidence that, in addition to the H1 helix and the L6 loop, the POTRA2 domain also undergoes conformational changes during secretion (Guerin et al., 2015). To determine which POTRA2 regions must be mobile for secretion, we searched for specific H bond- or salt bridge-mediated interactions present in the resting conformation (i.e., corresponding to the crystallographic structure) and disrupted them to loosen the structure or conversely replaced them with disulfide (S-S) bonds to limit motions of the corresponding regions. Of note, FhaC is naturally devoid of Cys residues. Residues involved in interactions between the POTRA2 domain and the barrel (Asn^245^-Ser^157^ and Asn^245^-Lys^184^) and in a barrel-distal region of the POTRA2 domain (Asp^165^-Lys^171^) were replaced with Ala or Cys, and the effects of these mutations on secretion activity were determined (Figure 2A-D). S-S bond formation is catalyzed by the periplasmic disulfide oxidase DsbA in the course of biogenesis, which generally affects SDS-PAGE migration of the protein in the absence of a reducing agent, unless the intervening loop between the Cys residues is too short. The Asn^245^Ala substitution markedly decreased the activity of FhaC and somewhat reduced its amount in the membrane, unlike formation of Cys^157^-Cys^245^ or Cys^184^-Cys^245^ S-S bonds (Figure 2C, D). This indicates that these barrel-POTRA2 interactions contribute to FhaC activity, possibly because they stabilize its conformation in the secretion cycle. On the contrary, the engineered Cys^165^+Cys^171^ substitutions strongly reduced the level and the activity of FhaC in a *dsbA^+^* background. Although protein migration was not affected, the S-S bond was most likely formed, since secretion was not reduced in a *dsbA^-^* background or with the individual substitutions. The observation that S-S bond formation between these two Cys residues is detrimental points to the need for flexibility in the barrel-distal region of the POTRA2 domain.

**Figure 2.**
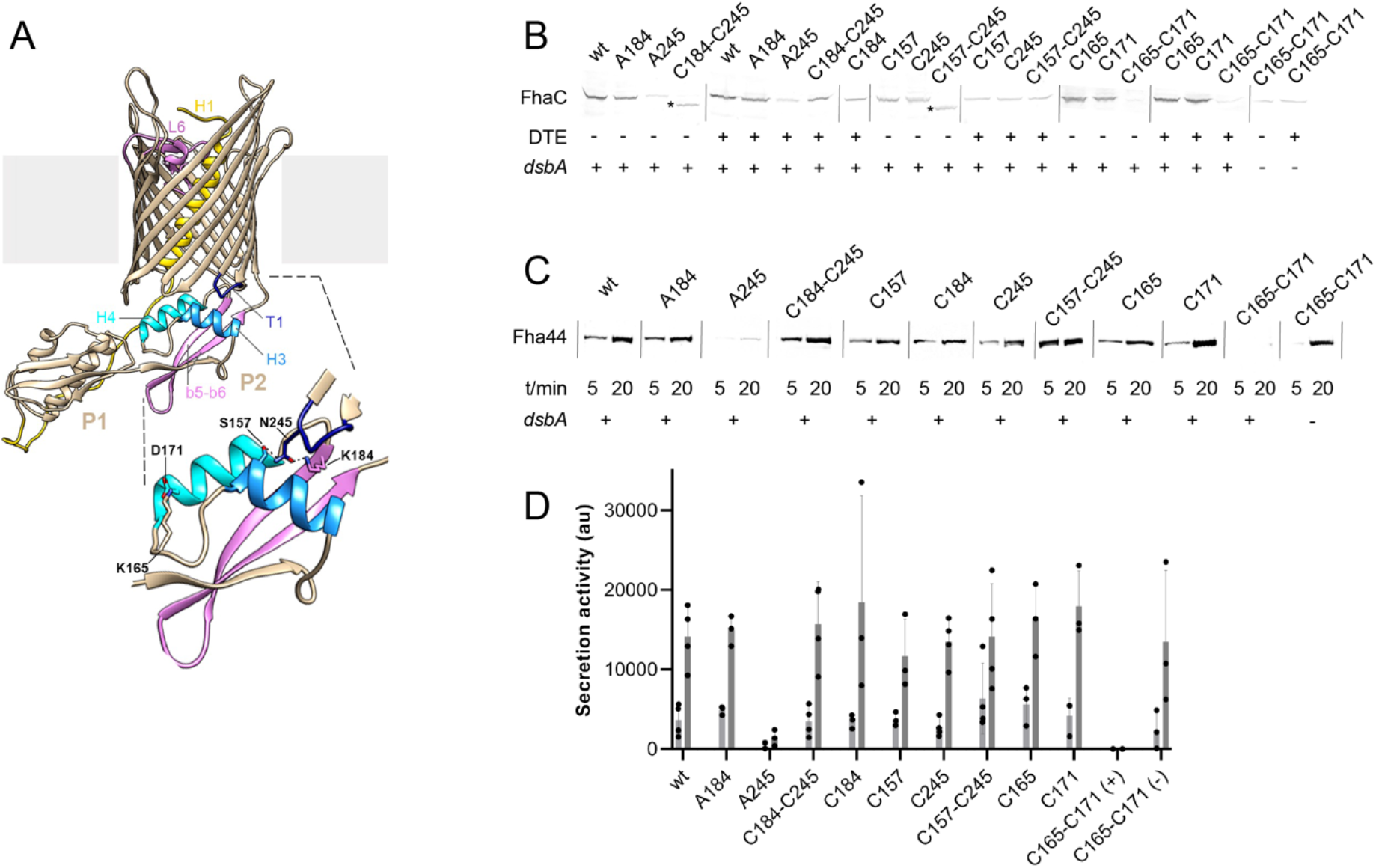
Effects of engineered S-S bonds on FhaC activity. (A) Structural model of FhaC (PDB 4QKY). A zoom of the POTRA2 domain is shown below. (B) Residues involved in a salt bridge (Lys^165^-Asp^171^) or H bonds (Lys^184^-Asn^245^; Ser^157^-Asn^245^) in the resting conformation of FhaC were replaced as indicated (C=Cys; A=Ala). Immunoblots were performed on membrane extracts with anti-FhaC antibodies. The asterisks indicate oxidized species of FhaC detected in the absence of the reducing agent dithioerythritol (DTE) in the sample buffer. (C) The secretion activity of the FhaC variants was determined using a model substrate, Fha44-His, affinity precipitated from supernatants 5 and 20 min after induction. Immunoblots were developed with an anti-6His tag monoclonal antibody. (D) Quantification of Fha44 found in culture supernatants after 5 and 20 min. The means and standard deviations of the means are shown (n=3 or 4). Activity of FhaC^C165+C171^ could only be detected in the *dsbA* KO strain (denoted C165-C171(-)), not in its wild type parent (denoted C165-C171(+)).

### Evidence for dynamics and alternative conformations of FhaC in lipid bilayers by NMR spectroscopy

To gain insight into the nature and the time scale of the conformational changes of FhaC, we made use of nuclear magnetic resonance (NMR) spectroscopy for its ability to characterize molecular structure and dynamics as well as minor conformational states of proteins in lipid bilayer environments (Mittermaier & Kay, 2009; Liang & Tamm, 2016). We recorded NMR spectra of FhaC in liposomes and lipid nanodiscs (Bayburt et al., 1998; Viegas et al., 2016), using solid- and solution-state NMR techniques, respectively. To render the 61-kDa protein more accessible to NMR spectroscopy, we resorted to perdeuteration and specific ^1^H, ^13^C-isotope labeling of isoleucine (Ile) δ_1_ methyl groups (Ruschak & Kay, 2010). Since the 15 Ile residues of FhaC are well distributed across all structural elements of the protein (Figure 3A), we expected this reduced labeling scheme to nevertheless be able to report on larger-scale structural transitions of FhaC.

**Figure 3.**
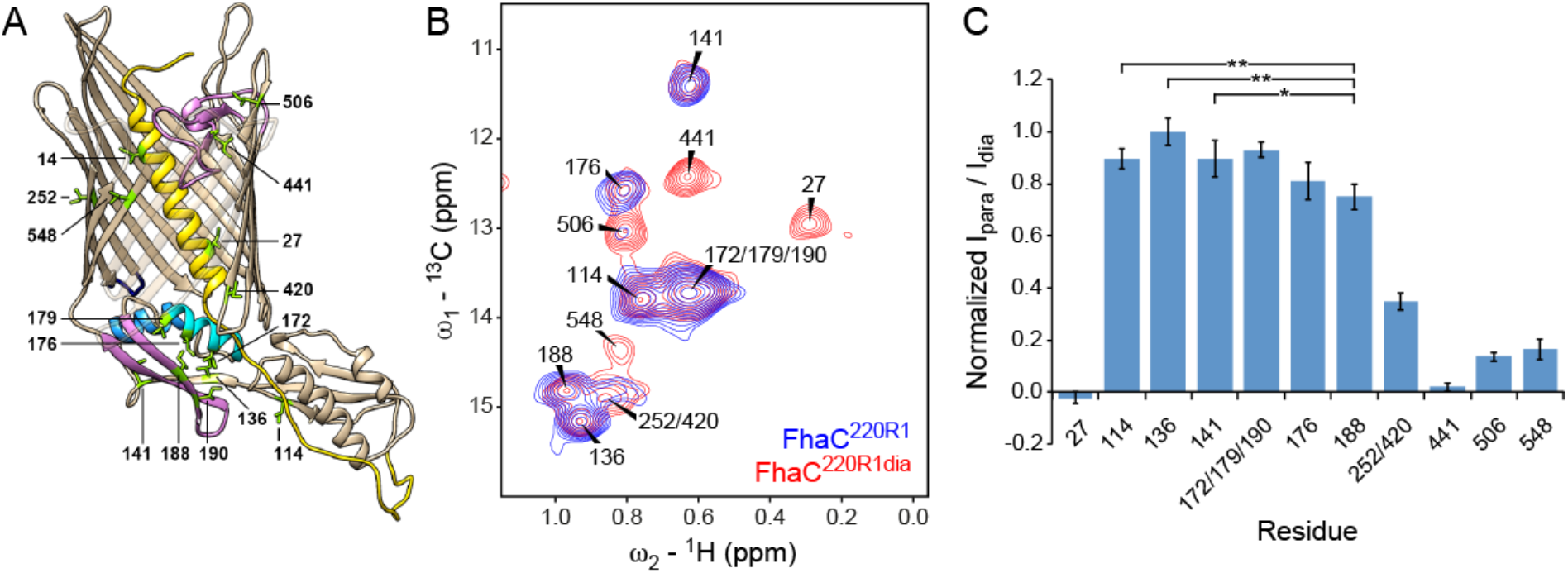
NMR analysis of Ile δ_1_-methyl labeled FhaC in lipid bilayers. (A) Structure of FhaC with Ile residues labeled and drawn as green sticks. Color code of structural elements is as in Fig. 1. β-strands 1 to 4 are drawn transparently for visibility. (B) Superposition of the methyl regions of solid-state dipolar hCH ^13^C-^1^H correlation spectra of u-(^2^H, ^15^N), Ile- δ_1_(^13^CH_3_)-labeled FhaC samples in *E. coli* polar lipid liposomes, with a paramagnetic MTSL tag (FhaC^220R1^, blue) or with a diamagnetic MTSL analog (FhaC^220R1dia^, red) attached to the introduced Cys^220^ residue. Spectra were recorded at 800 MHz ^1^H Larmor frequency and 50 kHz MAS. (C) Ratios *I*para/*I*dia of Ile-δ1 methyl peak intensities in the hCH correlation spectra of FhaC^220R1^ and FhaC^220R1dia^ shown in (B), normalized to the maximum ratio observed in Ile^136^. Error bars are calculated based on spectral noise levels. * and ** indicate significant (p < 0.05 and p < 0.01, respectively) attenuation of the Ile^188^ signal relative to the signals of the reference residues Ile^114^, Ile^136^, and Ile^141^. These analyses were complemented by analyses of scalar coupling-based spectra in liposomes and nanodiscs (Supplement 1), through-space correlations (Supplement 2), and relaxation dispersion experiments (Supplement 3). Estimated distances between MTSL probe and Ile residues in FhaC^220R1^ as expected from the crystal structure are shown in Supplement 4.

Signals from all Ile residues could be identified and assigned by Ile-to-Val mutations or paramagnetic relaxation enhancement experiments (Fig. 3B; see also below) (Amero et al., 2011; Venditti et al., 2011). The higher resolution of solution-state NMR spectra of FhaC in nanodiscs proved useful in the assignment (Figure 3 Supplement 1). Variable intensities of the Ile δ_1_ methyl signals report on local dynamics in the protein. While Ile^14^ in H1 was only visible in scalar coupling-based spectra, the signal of Ile^548^ in β-strand B16 at the barrel seam consistently exhibited low intensity in both scalar and dipolar coupling-based spectra (Figure 3B, Figure 3 Supplement 1). This indicates sub-µs time scale motion towards the N-terminus of the H1 helix and µs-to-ms time scale exchange dynamics at the barrel seam, respectively. The notion of dynamics in FhaC is also supported by the absence of through-space correlations for all but the shortest Ile-Ile distances expected from the crystal structure (Figure 3 Supplement 2). However, ^13^C rotating-frame (R_1ρ_) relaxation dispersion experiments probing exchange between states with different chemical shifts on the µs time scale (Lewandowski et al., 2011; Ma et al., 2014) yielded statistically flat dispersion profiles (Figure 3 Supplement 3), indicating that conformational changes of FhaC detectable by Ile δ_1_ methyl chemical shifts must occur on slower time scales.

To specifically probe for alternative FhaC conformations, we performed paramagnetic relaxation enhancement (PRE) NMR experiments in which a paramagnetic methanethiosulfonate spin label (MTSL) is attached to an engineered Cys in the protein, and attenuation of NMR signals of nuclei within a radius of about 25 Å around the MTSL probe can be detected even if they only transiently approach the probe (Battiste & Wagner, 2000; Clore & Iwahara, 2009; Nadaud et al., 2007). We chose residue 220 in the extracellular loop L1 for this experiment, yielding FhaC^220R1^, where R1 represents the spin label. Intensities of Ile δ_1_ methyl signals in FhaC^220R1^ measured by solid-state NMR in proteoliposomes were referenced to those in a sample with a diamagnetic MTSL analog attached to the same residue, FhaC^220R1dia^ (Fig. 3B, C). Comparison of the signal intensity ratios obtained for different FhaC residues allows to determine whether a signal is more attenuated than would be expected from the crystal structure, indicating a residue approaching the probe more closely (see Methods).

In agreement with expectations, residues more than 35 Å away from the position of the paramagnetic MTSL tag modelled onto the FhaC crystal structure (Ile^114^, Ile^136^, and Ile^141^ in POTRA1 and POTRA2) exhibited the highest para- versus diamagnetic intensity ratios, while residues expected to be within 16 to 25 Å of the paramagnetic center (Ile^27^, Ile^441^, Ile^506^, Ile^548^) showed attenuation of their NMR signals in paramagnetic FhaC^220R1^ (Figure 3B, C, Figure 3 Supplement 4). The overlapped signal corresponding to residues Ile^252^ and Ile^420^, at expected distances to the paramagnetic center of 12 and 32 Å, respectively, exhibited intermediate attenuation as expected. Signals from the POTRA2 H4 helix (Ile^172^, Ile^176^, Ile^179^) were not significantly attenuated compared to reference signals. However, the signal of Ile^188^ in strand b5 of the POTRA2 domain was attenuated more than would be expected for a residue at 35 Å distance from the paramagnetic center. The difference in attenuation with respect to the reference residues Ile^114^, Ile^136^, and Ile^141^ is significant (p < 0.05, Figure 3C). This result indicates that a region of the POTRA2 domain encompassing strand b5 can approach the extracellular loops of FhaC.

### Evidence for motions of the POTRA2 domain towards the extracellular side from EPR spectroscopy

To complement the NMR data, we resorted to electron paramagnetic resonance (EPR) spectroscopy, another technique suitable to detect dynamics and minor conformational states of proteins in lipid bilayers, but sensitive to longer distances than can be measured by NMR (Sahu & Lorigan, 2020; Torricella et al., 2021). Distances from about 1.8 to 8 nm between paramagnetic spin labels attached to membrane proteins can be measured with pulsed electron double resonance (PELDOR) EPR experiments and can provide insight into non-homogeneous conformational ensembles (Jeschke, 2012). Notably, in continuous-wave (CW) EPR spectroscopy experiments with FhaC carrying a single paramagnetic spin label attached at the solvent-exposed position 195 in the POTRA2 domain, we have previously observed a very slow-motion component for the spin probe (Guerin et al., 2015), suggesting possible interactions of the probe with the lipid bilayer in some conformers. To explore this further with explicit distance measurements, we performed PELDOR experiments. We introduced a Cys residue at position 503 in the extracellular L7 loop and combined it with another Cys either at position 195 in the b5-b6 hairpin of the POTRA2 domain, position 187 in the b5 strand, or position 33 in the linker, and we labeled both with an MTSL spin label, yielding FhaC variants FhaC^33R1+503R1^, FhaC^187R1+503R1^, and FhaC^195R1+503R1^.

In b-octyl glucoside (bOG) micelles, for FhaC^195R1+503R1^ and FhaC^187R1+503R1^, the main populated states correspond to distance distributions between the two spin probes that are consistent with distances calculated using MTSL rotamer libraries attached to the corresponding residues of the FhaC crystal structure (Figure 4A; Figure 4 Supplement 1) (Jeschke, 2013, 2020). For FhaC^33R1+503R1^, a broad distance distribution was observed, with contributions centered at 4.2 nm and 4.6 nm as predicted by rotamer libraries (Figure 4 Supplement 1). In proteoliposomes, for FhaC^195R1+503R1^ and FhaC^187R1+503R1^, the main populated states correspond to long distances of 5 to 6 nm between the two spin probes (Figure 4B). Note that these distances are shorter than the expected distances calculated using MTSL rotamer libraries, since the lipid environment limited the dipolar evolution times that could be applied in PELDOR experiments (Figure 4 Supplement 2). In addition to the expected long inter-spin distance, shorter distance distributions centered at 2.5 nm and 3.5 nm were observed for FhaC^195R1+503R1^ (Figure 4B). They can be attributed to conformers with the two spin probes closer to one another than in the crystal structure conformation. Similarly, additional peaks corresponding to shorter-than-expected distances were observed in the distance distributions for FhaC^187R1+503R1^ (Figure 4B). These results strongly support the idea that the b5-b6 hairpin of the POTRA2 domain moves towards the extracellular surface of FhaC in some conformers. For FhaC^33R1+503R1^, distances both shorter and longer than expected were obtained. This indicates that, in addition to moving away from the membrane when H1 exits from the pore, as reported in (Guerin et al., 2014), the linker also moves toward the surface in specific conformers.

**Figure 4.**
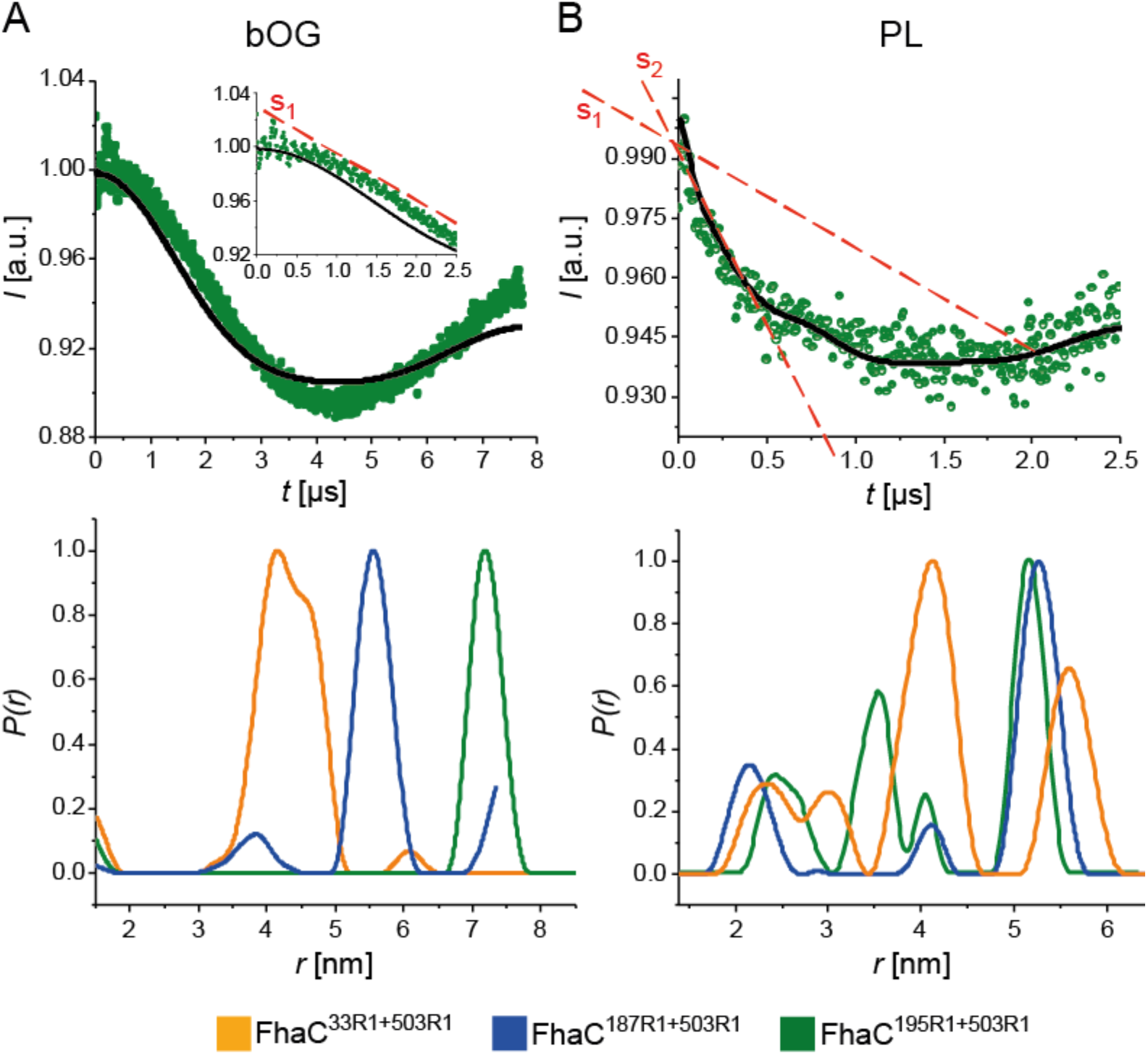
PELDOR analyses of FhaC. Dipolar evolution function for FhaC^195R1+503R1^ (top), and distance distributions obtained by Tikhonov regularization of the dipolar evolution functions (bottom) for FhaC^33R1+503R1^(orange), FhaC^187R1+503R1^ (blue), and FhaC^195R1+503R1^ (green) in bOG (A) and in proteoliposomes (B) prepared with *E. coli* polar lipids (PL). The black lines in the upper panels correspond to the fitting of the experimental PELDOR traces. The inset in A represents the first 2.5 µs of the dipolar evolution function for comparison with that shown in (B). The red dashed lines denoted S1 and S2 show the slopes of the first parts of the curves representing the dipolar evolution functions. Predicted distance distributions can be found in Supplement 1. Note that the longest distances measured depend on the dipolar evolution time *t*. As the lipid environment decreases the *t* that can be applied, the longest distances shift to smaller values for FhaC in proteoliposomes compared to bOG (Supplement 2). A mutation that severs the connection of the loop L6 to the inner barrel wall affects the EPR spectra of FhaC^195R1+503R1^ (Supplement 3). Spectroscopic analyses (PELDOR EPR and PRE NMR) of FhaC in nanodiscs complement these data (Supplement 4).

The point mutation Asp^492^Arg disrupts a conserved salt bridge between L6 and the inner barrel wall and induces conformational changes in FhaC (Guerin et al., 2015). To determine whether it affects the conformational equilibrium of the POTRA2 domain, we introduced the Asp^492^Arg substitution in FhaC^195R1+503R1^. Indeed, PELDOR experiments showed an increased proportion of species characterized by short inter-spin distances in this mutant (Figure 4 Supplement 3).

Notably, we did not obtain indications for alternative conformers in FhaC reconstituted into nanodiscs with EPR or NMR spectroscopy. Analysis of PELDOR experiments on FhaC^195R1+503R1^ in nanodiscs yielded only a long distance between the spin labels, as expected from the crystal structure, and PRE experiments on FhaC^195R1^ in nanodiscs showed attenuation of NMR signals only within the POTRA2 domain (Figure 4 Supplement 4). Along with smaller linewidths of FhaC NMR signals in nanodiscs compared to liposomes (Figure 3 Supplement 1), these findings indicate that the constrained nanodisc environment limits the conformational space accessible to FhaC and hinders the larger-scale conformational changes that can be observed in proteoliposomes.

Taken together, both our EPR and NMR data show that the POTRA2 domain can undergo large conformational changes that bring its b5-b6 hairpin close to the membrane and the extracellular side, and that these conformational changes are facilitated by the rupture of the interaction between L6 and the inner barrel wall. The H1-POTRA1 linker can also adopt alternative conformations and notably move towards the cell surface.

### *In vivo* evidence for conformers with the POTRA2 domain or the linker close to surface loops

To investigate whether the conformers observed in proteoliposomes also exist *in vivo* and to obtain insight into the potential position of the POTRA2 domain in those conformers, we simultaneously replaced two residues distant in the X-ray structure of FhaC with Cys residues to detect spontaneous S-S bond formation. Our rationale was that conformational changes that bring the two Cys residues close to each other should promote S-S bond formation even if the corresponding alternative conformations are short-lived, as the S-S bound species accumulate over time. These experiments were performed in a *dsbA^-^* background, such that S-S bonds formed after FhaC biogenesis and thus exclusively resulted from its conformational changes in the membrane. We combined Cys residues at the extracellular surface at positions 224 in L1, 290 in L3, 342 in L4, 391 in L5, 503 in L7 or 545 in L8 with periplasmic Cys residues in the POTRA2 domain at positions 167, 176 or 195, in the linker at position 48, or in the POTRA1 domain at position 86 (Figure 5A). None of the single Cys substitutions markedly affected the secretion activity of FhaC (Baud et al., 2014; Guerin et al., 2014; Guerin et al., 2015). Under non-reducing conditions, partial oxidation of FhaC as detected by aberrant migration in SDS-PAGE was identified for the combinations Cys^195^+Cys^224^, Cys^176^+Cys^224^, Cys^48^+Cys^224^, Cys^48^+Cys^545^ and weakly for Cys^167^+Cys^224^, indicating S-S bond formation within specific pairs of engineered Cys residues (Figure 5B). In contrast, no loop other than L1 or L8 was found to cross-link with those periplasmic regions, and none cross-linked with the Cys residue in the POTRA1 domain. To confirm S-S bond formation, the FhaC^C48+C224^ and FhaC^C195+C224^ variants were overexpressed, purified and subjected to liquid chromatography coupled to tandem mass spectrometry (MS) in reducing and non- reducing conditions. In both variant samples, the regions that contain the Cys residues were detected only when proteolytic digestion was performed after reduction and alkylation (Figure 5 Supplement 1), which supports S-S bond formation between the linker and L1 in FhaC^C48+C224^ and between the POTRA2 domain and L1 in FhaC^C195+C224^. Thus, *in vivo*, the last portion of the linker can be found close to the extracellular loops L1 and L8 that immediately follow and precede the first and last β−barrel strands, B1 and B16, respectively, and the α helix H4 and the b5-b6 β hairpin of the POTRA2 domain can be found close to the extracellular loop L1. This indicates that these periplasmic elements approach the β−barrel seam in specific conformers, in agreement with our *in vitro* data.

**Figure 5.**
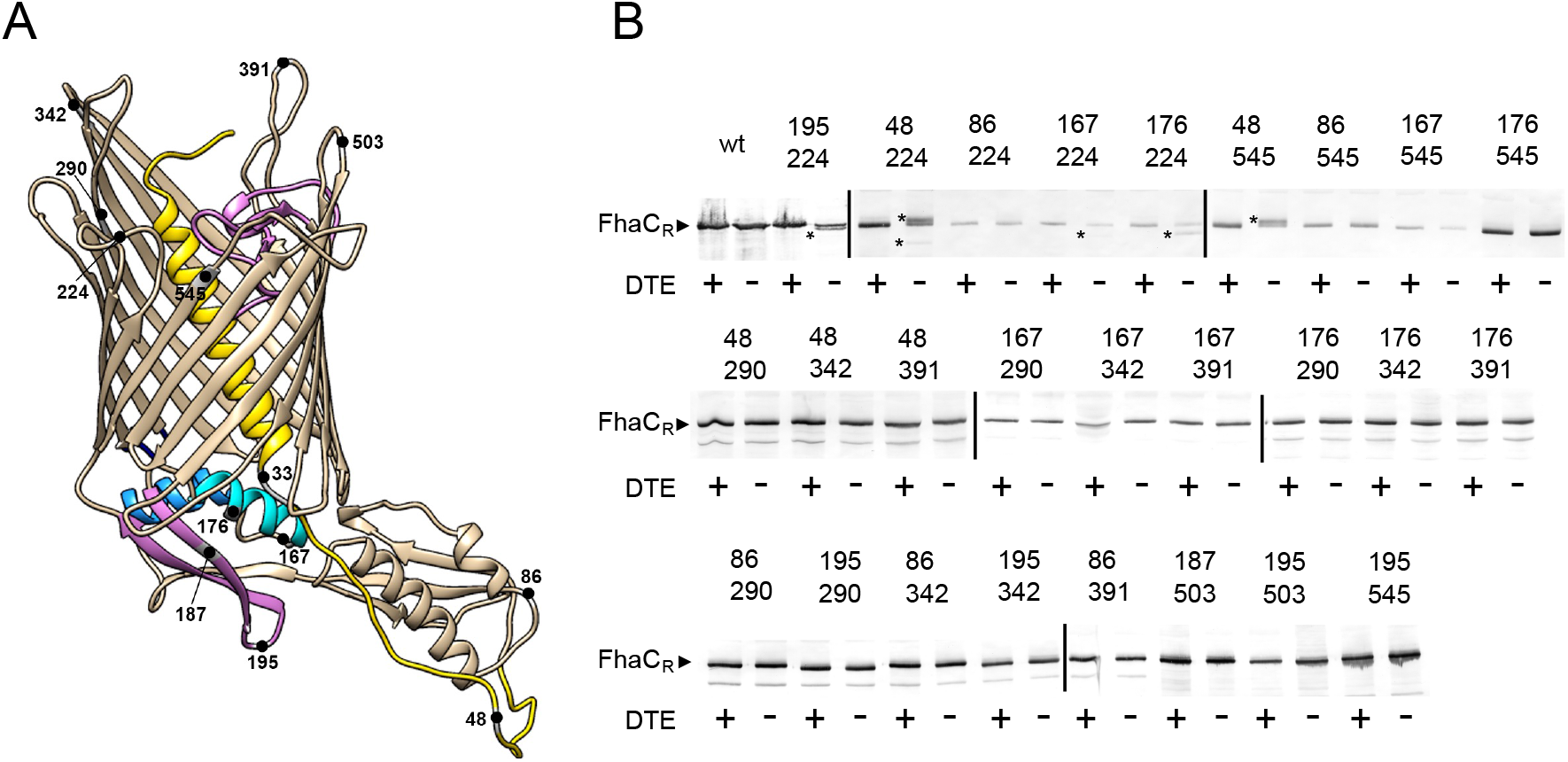
Detection of transient conformers of FhaC *in vivo*. (A) Position of the Cys substitutions in FhaC. (B) Immunoblot of membrane fractions of *E. coli JCB571* (*dsbA* KO strain) producing FhaC variants. The numbers indicate the positions of the two Cys residues. The reducing agent DTE was added to one half of each sample. FhaC_R_ represents the position of the reduced form. The asterisks point to the additional, cross-linked forms that can migrate faster or more slowly than the reduced form, depending on the respective positions of the two Cys residues. S-S bond formation was confirmed by mass fingerprinting analyses (Figure Supplement 1).

### Interactions of portions of the POTRA2 domain with the β barrel by native mass spectrometry

Our data imply that the POTRA2 domain undergoes some breaking up in the secretion cycle. We thus investigated its lability by using structural MS-based approaches (Figure 6). Native MS analysis of FhaC revealed a monomer that could be stripped of bOG at 60 V, a relatively low collision energy (CE), and that displayed a narrow charge state distribution between 14+ and 19+ indicative of a folded protein in a single conformation (Figure 6 A, B, Figure 6 Supplement 1).

**Figure 6.**
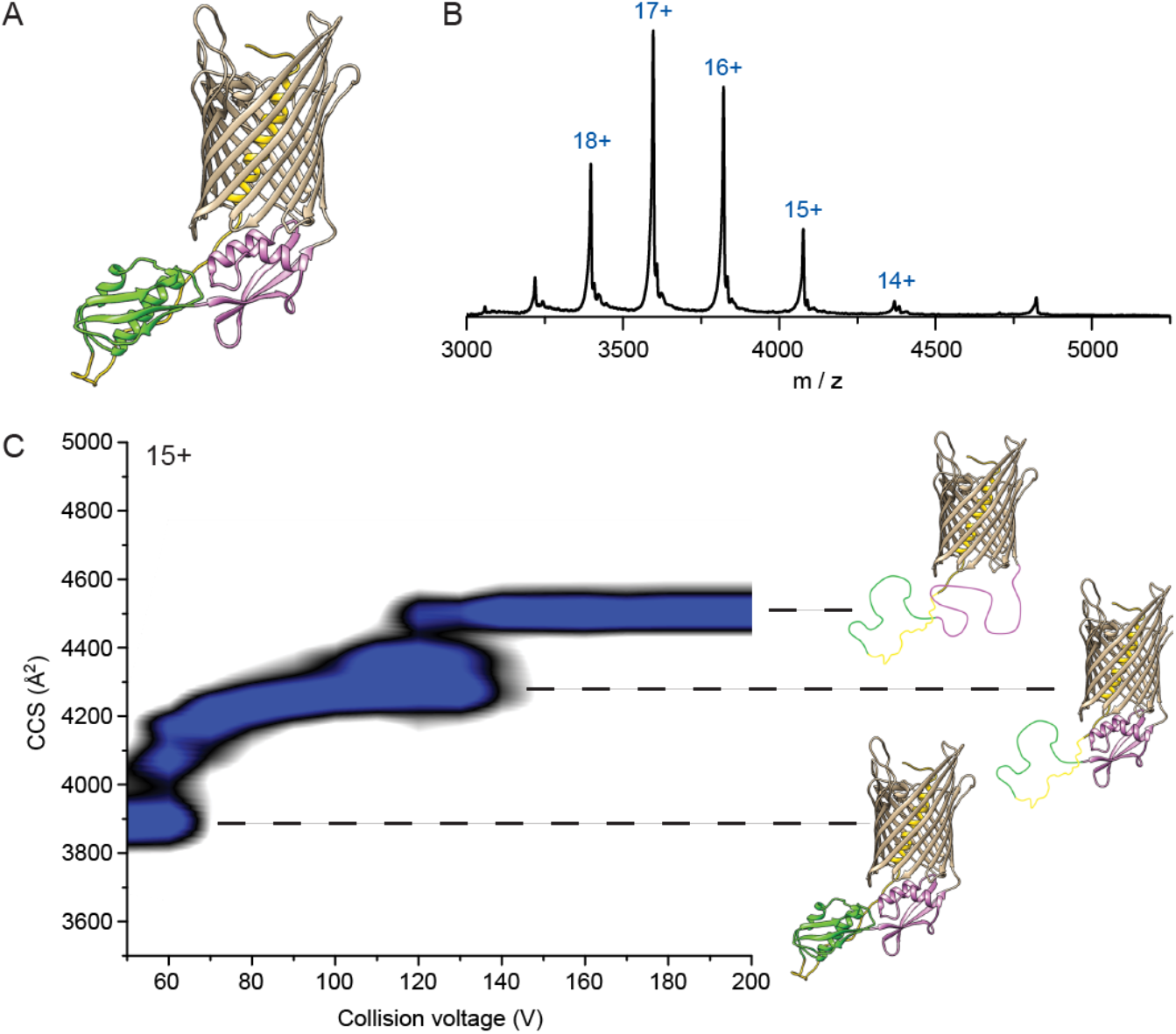
Native mass spectrometry analysis of WT FhaC. (A) Model of FhaC with H1 and the linker in yellow, and the POTRA domains 1 and 2 in pink and green, respectively. (B) Mass spectrum of WT FhaC released from its bOG micelle. The spectra at increasing collision energy are shown in Supplement 1. (C) Collision-induced unfolding (CIU) experiments show two dominant transitions that are likely linked to unfolding of the POTRA domains (see text), although the order in which they unfold is unknown. CIU profiles of control *β*-barrel proteins are shown in Supplement 2, and profiles of the FhaC^C4+C391^ variant with H1 locked in the barrel are in Supplement 3.

We used collision-induced unfolding (CIU) (Tian et al., 2015) to characterize the stability and the organization of the FhaC domains. FhaC displayed two transitions at 60 V and 120 V as shown by the increases of collision cross section (CCS) values (Figure 6C). As the number of transitions in the gas phase can generally be related to the number of domains of a protein (Zhong et al., 2014), and extra-membrane domains are more likely to experience early unfolding than domains embedded in detergent or lipids due to collisional cooling (Barrera et al., 2009), those transitions might be caused by unfolding of the POTRA domains and/or ejection and unfolding of H1. Control CIU experiments with other *B. pertussis* outer membrane proteins (OMPs) with small soluble domains inside their β barrels, the TonB-dependent transporter BfrG and the translocator domain of an autotransporter, SphB1-αβ, showed a single unfolding transition at low voltage, which likely corresponds to unfolding of these soluble domains (Figure 6 Supplement 2). Thus, the β barrels of these three proteins likely remain structurally intact at high activation conditions, most likely due to strong hydrogen bonding networks.

To further investigate whether the CIU transitions observed for FhaC stem from unfolding of the POTRA domains or ejection of the H1 helix, we studied the CIU pathway of FhaC^C4+C391^ in which H1 is locked inside the barrel by an S-S bond and thus cannot move out (Guerin et al., 2014). FhaC^C4+C391^ exhibited the same transitions as wt FhaC, although the second unfolding event was delayed by 30 V and the overall CCS value was increased by 50 Å^2^ (Figure 6 Supplement 3). As CIU is unlikely to break S-S bonds (Tian et al., 2015), comparison of these unfolding pathways suggests that the two transitions correspond to successive unfolding of the POTRA domains, with the barrel remaining intact in those conditions. H1 stays inside the barrel or its unfolding barely registers in the CCS values. The delay of the second unfolding transition for FhaC^C4+C391^ suggests that locking H1 in the barrel stabilizes one of the POTRA domains, although from the data we cannot discern which one.

We next tested the possibility that portions of the POTRA2 domain might bind to the β barrel, probably along strands B1 or B16 upon opening of the barrel seam. Using native MS, we assessed the binding of synthetic peptides that correspond to various periplasmic portions of FhaC, including b5-b6, b4+L (*i.e.*, b4 followed by the b4-H3 linker) and L+H4 (*i.e.*, the H3-H4 linker followed by H4) of the POTRA2 domain, b2-b3 of the POTRA1 domain, Lk, a non-structured peptide from the linker region between H1 and the POTRA1 domain, and the N-terminal β hairpin of the FhaB transport substrate, Fha-NT (Figure 7; Figure 7 Supplement 1). The same experiments were performed with SphB1-αβ to correct for non-specific binding, which might occur in native MS experiments due to artifacts induced by interaction with the detergent during the electrospray process (Landreh et al., 2016). Fha-NT, b4+L and b5-b6 exhibited binding to FhaC, with b5-b6 binding at the highest level and in two copies, but markedly less to the FhaC^C4+C391^ variant (Figure 7A; Figure 7 Supplement 2). In contrast, the peptides containing the sequences of b2-b3 of the POTRA1 domain, H4 or the H1-POTRA1 linker did not bind.

**Figure 7.**
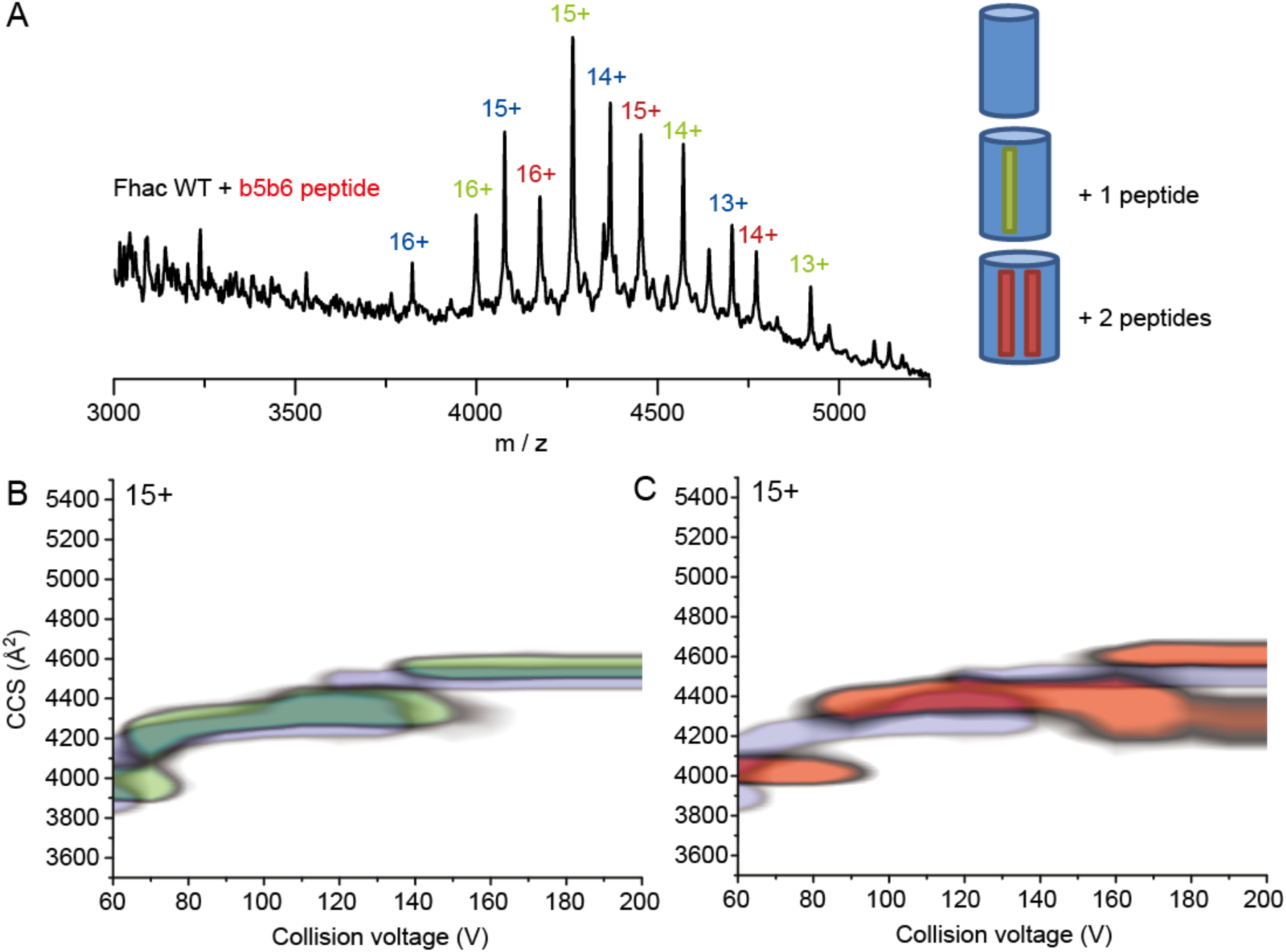
Binding of synthetic peptides to FhaC. (A) Mass spectrum of FhaC incubated with the b5-b6 peptide at a collisional energy of 150 V shows binding of the peptide to the protein under high-energy conditions. Schematic representations of the barrel with bound peptides are shown at the right. (B) Overlay of the CIU plots of FhaC with (green) and without (blue) one b5-b6 peptide bound. Increased CCS values are observed both under native conditions (low CE) and conditions in which the POTRA domains are most likely unfolded (high CE). (C) Comparison of the CIU plots of FhaC alone (blue) and FhaC with two b5-b6 peptides bound (orange), which shows an additional CCS increase compared to FhaC with a single peptide bound. The peptides used in this study are shown in Supplement 1. Binding of a different peptide and quantification of the binding data are shown in Supplement 2, the CIU plots of wt FhaC with the b4+L and Fha-NT peptides are shown in Supplement 3, and the CCS values measured in those experiments are in Supplement 4. The CIU plot of FhaC^C4+C391^ with the b5-b6 peptide is shown in Figure 6 Supplement 3.

We assessed structural changes induced by peptide binding using native ion-mobility (IM) MS. At low CE (*i.e.*, no activation), all three peptides increased the CCS of the compact state of FhaC by rather small increments of 91-92 Å^2^ (Figure 7B, C; Figure 7 Supplement 3). However, upon increasing the activation conditions, Fha-NT and b4+L no longer increased the CCS of FhaC, compared to the unbound protein. In contrast, the b5-b6 peptide caused an increase in CCS values both at low and high collisional activation, suggesting that a structural change was induced upon peptide binding and that the peptide was bound to a region that remains folded in these conditions (Figure 7B, C, Figure 7 Supplement 4). As our CIU studies suggest that the POTRA domains likely unfold at high CE, the effect of b5-b6 on the CCS might thus stem from peptide binding to the β barrel. The same experiment with FhaC^C4+C391^ showed a lower level of peptide binding, which nevertheless caused a similar increase of CCS at both low and high energies, like with wt FhaC (Figure 6 Supplement 3, Figure 7 Supplement 4). This supports the model that the peptide corresponding to the b5-b6 hairpin of the POTRA2 domain interacts with the β barrel, and that this interaction is facilitated by the ejection of H1. Given the amphipathic nature of this hairpin, it most likely aligns with an edge of the open β-barrel seam, consistent with the cross-linking data.

## DISCUSSION

As Omp85 transporters are thought to perform their functions in the absence of an energy source in the periplasm, their postulated conformational cycling must involve low energy barriers between conformers, as reported for BamA (Xiao et al., 2021). Here, we obtained evidence for large conformational changes of FhaC that involve portions of the POTRA2 domain approaching the extracellular side of the protein. Conformational changes of FhaC occur independently of the presence of the substrate, indicating that such dynamics is an intrinsic structural feature of the protein, with implications for its function. Notably, the conformational states appear to be in slow equilibrium, as with BamA (Hartmann et al., 2018).

All structural elements of TpsB transporters are connected with one another, structurally and functionally, and their motions appear to be coupled (Guerin et al., 2014; Guerin et al., 2020; Guerin et al., 2015; Maier et al., 2015). In the resting conformation, H1 and L6 interact with the barrel wall, H1 with L1, the H1-POTRA1 linker with the POTRA domains, and the POTRA2 domain with the periplasmic side of the barrel (Guerin et al., 2020; Maier et al., 2015). In the secretion process, L6 breaks its connection with the barrel wall, H1 moves towards the periplasm, and part of the B1-B16 seam unzips (Guerin et al., 2014; Guerin et al., 2020; Guerin et al., 2015). In the current work we have obtained evidence fur further conformational changes involving the POTRA2 domain. Thus, EPR, NMR and S-S cross-linking data revealed the proximity of parts of the POTRA2 domain to the barrel seam and to the extracellular side of FhaC in specific conformers, and structural MS experiments showed the binding of the POTRA2 b5-b6 hairpin peptide to the β barrel under conditions in which the POTRA domains are very likely unfolded.

Based on these and previous data, we propose the following model for the first steps of secretion (Figure 8). The closed, resting conformation of FhaC is in slow equilibrium with an open conformation in which L6 has been released from its interaction with the barrel wall and H1 has moved out of the pore (Figure 8A) (Guerin et al., 2014; Guerin et al., 2020; Guerin et al., 2015). In the open conformation, the linker has vacated the substrate binding site on the POTRA2 domain, thus enabling a specific portion of the conserved TPS domain of the substrate to bind to the groove between H4 and b5 (Delattre et al., 2011). According to molecular dynamics simulations, the exit of H1 facilitates barrel unzipping between B1 and B16 (Guerin et al., 2020). Barrel unzipping is coupled with a swing motion of the b5-b6 hairpin of the POTRA2 domain towards the barrel, followed by its binding to the open seam, probably by β augmentation, thus enlarging the barrel and hoisting the bound substrate into the channel (Figure 8B).

**Figure 8.**
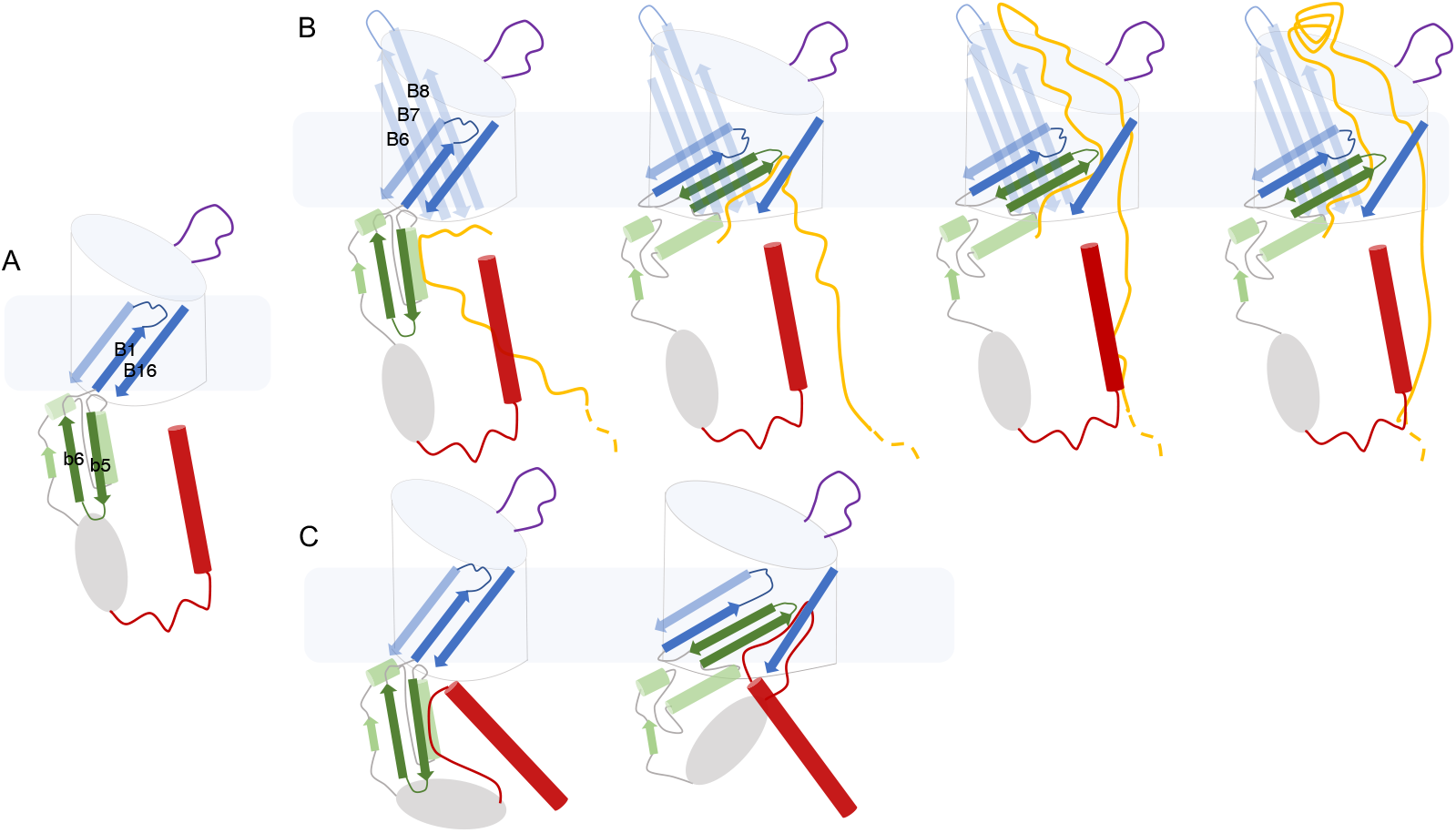
Model for TPS secretion. In the open conformation of FhaC (A), the substrate binding groove between H4 and b5 in the POTRA2 domain is available. (B) A specific region of the TPS domain of the substrate (yellow) binds to the POTRA2 domain, likely by β augmentation of the b5 strand. Unzipping of the barrel seam and insertion of the b5-b6 hairpin hoist a first portion of the substrate into the channel. The substrate likely forms a hairpin inside the barrel. Its diffusion toward the surface enables a specific portion of the TPS domain to interact with the B5-B8 beta sheet that protrudes at the cell surface. This interaction templates the folding of the substrate into a nascent β helix at the cell surface. The substrate is progressively threaded through the channel and folds at the surface, until a stable nucleus has formed. (C) In the absence of the substrate the linker binds to the POTRA2 domain, and the conformational changes of the protein can hoist it towards the cell surface in a futile cycle.

The next secretion steps are more speculative. The entropic cost of confining a portion of the bound, unfolded polypeptide inside the channel might facilitate its diffusion toward the surface (Halladin et al., 2021). The TPS domain most likely forms a hairpin inside the barrel, as proposed previously (Nash & Cotter, 2019), and a specific region binds to the extracellular β sheet formed by B5-B8 (Baud et al., 2014), which might template the folding of the substrate into a nascent β helix. The substrate would progressively be threaded across the channel and fold at the surface, leading to the formation of a stable β helix nucleus (Alsteens et al., 2013), which would prevent backtracking of the protein chain, effectively adding directionality to its stochastic motion as proposed by the “Brownian ratchet” mechanism (Peterson et al., 2010). According to our model, TpsB transporters mediate protein secretion without the need for cofactors by utilizing intrinsic protein dynamics. Although the conformational changes we describe are unprecedented in the Omp85 superfamily, divergent functional evolution has necessarily led to specific mechanistic adaptations. Recent work has indicated that energy may be transduced from the inner membrane to the BAM complex through the protonmotive force-utilizing SecDF complex (Alvira et al., 2020). One cannot rule out that the intrinsic conformational changes of FhaC *in vivo* are similarly enhanced by an energy-transducing mechanism.

The b5-b6 hairpin most likely binds by β augmentation of the barrel strand B1, by analogy with BamA and Sam50 in which the unzipped B1 strand templates folding of client proteins by β augmentation (Doyle & Bernstein, 2019; Höhr et al., 2018; Tomasek et al., 2020; Wu et al., 2021). This mode of binding is supported by our CIU results showing an increased cross-section of the protein upon binding of b5-b6 to the barrel and by the *in vivo* formation of an S-S bond between the tip of that hairpin and the extracellular loop L1. The b5-b6 sequence, with its amphipathic nature and suitable charge partitioning, fits ideally in the open seam. The observation that portions of the linker may approach the extracellular loops likely reflects futile conformational changes involving the linker bound to the POTRA2 being hoisted into the channel in place of the substrate, in the absence of the latter (Figure 8C). Notably, evidence for the C-terminal portion of the linker reaching the cell surface was obtained previously (Guedin et al., 2000).

As the POTRA2 domain partially breaks up during secretion, it must reassemble after secretion is completed. This may be mediated by the interactions of the H3 helix and the barrel-proximal end of b5 of the POTRA2 domain with the periplasmic turn T1 of the barrel, which are important for FhaC activity. These fixed points of the POTRA2 domain may ensure that FhaC can regain its resting conformation after secretion, which is necessary to limit outer membrane permeability. Consistent with this hypothesis, disrupting the conformation of the periplasmic turn T1 yielded transient, very large channels as detected in electrophysiology experiments (Méli et al., 2006). Stabilizing specific conformations of the transporter might also account for the importance of the interaction between L6 and the inner barrel wall (Delattre et al., 2010).

In summary, we propose a novel mechanism of protein transport in TPS systems based on large-scale, spontaneous conformational dynamics of the TpsB partner. Our model integrates and explains currently available data from several complementary *in vitro* and *in vivo* approaches and establishes mechanistic links between TpsBs and other Omp85 transporters.

## Materials and Methods

### Strains and plasmids

*E. coli* JCB570 or JCB571 (*dsbA*::kan) were used for low level expression of FhaC and *E. coli* BL21(DE3*-omp5*) for overexpression. For peptide mapping FhaC^C48+C224^ and FhaC^C195+C224^ were overexpressed in BL21(DE3*-omp5 dsbA::kan*), which was constructed as described in (Derbise et al., 2003). Point mutations in *fhaC* were generated using the QuikChange II XL Kit (Agilent, Les Ulis, France) on pFc3 (Guedin et al., 2000). Overexpression of FhaC for purification was performed from pET22 or pET24 plasmids (Clantin et al., 2007). ptacFha44-His codes for the first 80 kDa of the FhaC substrate FhaB, called Fha44, followed by a 6-His tag. It was constructed by adding a 1.2-kb Sal-BamHI fragment of the *fhaB* gene into the same sites of ptacNM2lk-His (Guerin et al., 2015). pFJD63 codes for FhaC under the control of the P_BAD_ promoter (Guedin et al., 1998). Its derivatives were constructed by ligating the XhoI-HindIII and XhoI-XbaI fragments of pFJD63 with the XbaI-HindIII *fhaC* fragments carrying the relevant mutations from the pFc3 derivatives. pMSP1D1 and pMSP1E3D1 were obtained from Addgene (Watertown, MA, USA). pSphB1αβ is a derivative of pT7SBαβ (Dé et al., 2008) with a 6-His tag. To construct pT7bfrG- H, the sequence corresponding to the mature protein was PCR amplified and inserted in pFJD138 (Méli et al., 2006) after the signal-peptide and 6-His tag sequences.

### In vivo assays

To monitor S-S bond formation *in vivo*, the pFc3 variants were introduced in *E. coli* JCB571. The recombinant bacteria were grown at 37°C in minimum M9 medium containing 0.1% casaminoacids under agitation. The cells were collected by centrifugation when the optical densities at 600 nm (OD_600_) of the cultures reached 0.8. The cell pellets were resuspended in 50 mM sodium (pH 6.8) containing 10 mM N-ethylmaleimide and lysed using a Hybaid ribolyzer apparatus (50 sec at speed 6). The membranes were collected by ultracentrifugation of the clarified lysates at 90,000 g for 1 h. The pellets were resuspended in loading buffer without reducing agent and separated into two aliquots, with DTE added at 25 mM to one of them before heating at 70°C for 10 min. FhaC was detected using anti-FhaC antibodies (Delattre et al., 2011) with alkaline phosphatase development for 15 min.

For the secretion assays, overnight cultures of *E. coli* JCB570 or JCB571 harboring a pFJD63 derivative and ptacFha44-His were diluted to OD_600_ of 0.3 in LB and grown under agitation with 0.01% arabinose for 20 min to produce FhaC. The bacteria were collected by centrifugation, resuspended in prewarmed LB without arabinose and grown to OD_600_ of 0.8 before adding IPTG at 1 mM to induce the expression of Fha44. Culture aliquots were collected 5 and 20 min thereafter and placed on ice. After centrifugation to harvest the bacteria, Fha44 was affinity-purified from the supernatants with Ni-NTA beads (Qiagen, Courtaboeuf, France). The membrane extracts were prepared and FhaC was detected as above. Fha44 was detected by immunoblotting using anti-6His antibodies, the ECL kit of Amersham (Merck, St Quentin-Fallavier, France) and the Amersham Imager 600 (GE) with 1 sec exposure. The amounts of Fha44 in supernatants were quantified with ImageJ.

### Protein Purification and spin labeling

The production and purification of the FhaC derivatives was performed as described (Guerin et al., 2014). Expression for NMR experiments was performed in M9 minimal medium in D_2_O, 2.5 g/L ^2^H-glucose (Sigma, St Quentin-Fallaviers, France), 1g/L ^15^N-NH4Cl, 1g/L ^15^N,^2^H-isogro (Sigma) and ^13^C-α-ketobutyric acid (Sigma) to achieve u-(^2^H,^15^N), Ile-δ_1_(^13^CH_3_) isotope labeling (Ruschak & Kay, 2010). For spin labeling, 3 mM tris(2-carboxyethyl)phosphine (TCEP, Sigma) was added to the detergent extract before ion exchange chromatography. The FhaC-containing fractions were mixed with a 10-fold molar excess (1-oxyl-2,2,5,5-tetramethyl-Δ3-pyrroline-3-methyl) methanethiosulfonate (MTSL) or its diamagnetic analogue (1-Acetoxy-2,2,5,5-tetramethyl-δ-3- pyrroline-3-methyl) methanethiosulfonate (Toronto Research Chemicals, North York, ON, Canada) at 15°C with gentle agitation for 16 hours. Excess MTSL was removed by chromatography. SphB1-αβ and BfrG were produced from *E. coli* BL21(DE3*-omp5)* and purified from bOG extracts using Ni^2+^ affinity chromatography. For BfrG 300 mM NaCl was added to improve solubility.

### Preparation of liposomes and nanodiscs and protein reconstitution

Small unilamellar vesicles (SUVs) of *E. coli* polar lipids were prepared as described (Guerin et al., 2014). The SUVs were mixed with FhaC variants at lipid:protein molar ratios of approx. 2500:1 for EPR and 200:1 for NMR experiments, respectively, at room temperature, with gentle agitation for one hour. The proteoliposomes were formed by removal of detergent with the progressive addition of Biobeads SM2 (Bio-Rad), and the proteoliposomes were collected by ultracentrifugation. All steps were performed under argon. Final buffer concentrations after mixing FhaC and liposomes were about 12.5 mM each of Tris-HCl and NaP_i_, 150 mM NaCl, pH 6.7.

Nanodiscs were prepared with the MSP1D1 and MSP1E3D1 scaffold proteins (Ritchie et al., 2009) produced in *E. coli* BL21(DE3), with an induction of 3 h at 28°C. For NMR experiments, scaffold proteins were expressed in M9 minimal medium in D_2_O using ^2^H-glucose as carbon source to suppress their signals in the (^1^H,^13^C)-based NMR spectra. The bacteria were broken using a French press in 50 mM Tris-HCl (pH 8), 300 mM NaCl (TN buffer), 1% Triton X100 (TNX buffer), and the clarified lysates were subjected to Ni^2+^ affinity chromatography. After successive washes in TNX, TN buffer with 50 mM cholate, 20 mM and 50 mM imidazole, the proteins were eluted in TN buffer with 400 mM imidazole, concentrated by ultrafiltration and dialyzed against 20 mM Tris-HCl (pH 8), 200 mM NaCl and 0.1 mM EDTA. Dimyristoyl phosphatidyl choline (DMPC) and dimyristoyl phosphatidyl glycerol (DMPG) (Avanti, Interchim, Montluçon, France) at a 2:1 ratio were solubilized in chloroform, lyophilized overnight and resuspended to 25 mM in 20 mM Tris-HCl (pH 7.5), 100 mM NaCl, 0.5 mM EDTA, 50 mM cholate. For NMR experiments, deuterated (d_54_-) DMPC and DMPG (Cortecnet, Voisins-le-Bretonneux, France) were used. FhaC, the scaffold protein and the lipids were mixed at a ratio of 1:3:180, cholate was added to 15 mM, and incubation was performed for 1 h at room temperature. Biobeads were added progressively, and the incubation was continued at 4°C overnight. The nanodiscs were collected by ultracentrifugation and concentrated by ultrafiltration. For NMR experiments, the buffer was exchanged to 100 mM NaP_i_ in D_2_O pH* 7.2 using a 2-ml ZebaSpin column (7 kDa MWCO).

### NMR experiments

For solid-state NMR experiments on FhaC variants reconstituted into liposomes, the proteoliposomes collected by ultracentrifugation were transferred to 1.3 mm magic-angle-spinning (MAS) solid-state NMR rotors (Bruker Biospin, Wissembourg, France) using an ultracentrifugation device (Bertini et al., 2012) (Giotto Biotech, Sesto Fiorentino, Italy) in a Beckman ultracentrifuge (SW 32 Ti rotor, 77,000 x g, 12°C, 30 – 60 min). NMR experiments were performed on spectrometers operating at 800 and 950 MHz ^1^H Larmor frequency (18.8 and 22.3 T magnetic field) (Bruker Biospin) at a MAS frequency of 50 kHz. Sample temperature was kept at about 17°C as judged by the chemical shift of the bulk water resonance. Spectra were indirectly referenced to 2,2-dimethyl-2-silapentane-5-sulfonate (DSS) via the lipid methylene proton resonance, which appears at 1.225 ppm under our experimental conditions. Typical pulse lengths for ^1^H and ^13^C hard 90° pulses were 2.1 and 3.8 µs, respectively. For cross-polarization (CP), field strengths were 21 and 30 kHz for ^1^H and ^13^C, respectively (n=1 double-quantum Hartmann-Hahn condition), with a 50-to-100% ramp on the ^1^H radiofrequency (RF) field and a duration of 1.5 ms. ^1^H-detected 2D ^13^C-^1^H dipolar hCH correlation spectra (Barbet-Massin et al., 2014) were typically recorded with 1600 data points and a spectral width of 40 ppm in the direct ^1^H dimension and 100 to 140 data points and a spectral width of 13 ppm in the indirect ^13^C dimension. For water suppression, the MISSISSIPPI scheme (Zhou & Rienstra, 2008) at 15 kHz ^1^H RF field with a duration of typically 200 ms was employed. For the 2D hChH correlation spectrum, a ^1^H-^1^H mixing time of 6.4 ms using radio frequency driven recoupling (Bennett et al., 1992) with a ^1^H field strength of 120 kHz was applied between back-CP and acquisition. ^13^C R1ρ spectra (Lewandowski et al., 2011; Ma et al., 2014) were recorded in a pseudo-3D fashion, with the ^13^C spinlock period inserted between the initial CP and the ^13^C indirect evolution of the hCH sequence. Spinlock field strengths from 1.2 to 10 kHz were used, and 5 spinlock durations from 2.5 to 80 ms with one repeated value were recorded for each spinlock. The spinlock carrier frequency was kept at the center of the isoleucine δ1 methyl ^13^C region, as in all other hCH correlation spectra. A ^1^H 180° pulse was inserted in the middle of the spinlock period to suppress chemical shift anisotropy / dipolar coupling cross-correlated relaxation (Kurauskas et al., 2016). Solid-state PRE NMR experiments were recorded on FhaC samples with either a paramagnetic MTSL tag or a diamagnetic MTSL analogue (Nadaud et al., 2007) attached to a Cys, reconstituted into *E. coli* polar lipid liposomes. Standard dipolar 2D hCH correlation spectra were recorded.

Solution-state NMR experiments on FhaC in nanodiscs were conducted on a 900 MHz spectrometer (Bruker Biospin) at 32°C sample temperature. Standard ^13^C-^1^H heteronuclear multiple-quantum coherence (HMQC) or SOFAST-HMQC (Schanda & Brutscher, 2005) experiments were recorded with 2048 and 150 data points and spectral widths of 14 and 7.4 ppm in direct ^1^H and indirect ^13^C dimensions, respectively. For PRE experiments, standard ^13^C-^1^H HMQC spectra were recorded on a FhaC^195R1^ sample before and after reduction of the paramagnetic MTSL tag with a 10-fold molar excess of ascorbic acid (Battiste & Wagner, 2000). NMR spectra were processed with TopSpin 4.0.3 (Bruker Biospin) or NMRPipe (Delaglio et al., 1995) and analyzed with Sparky (Lee et al., 2015) or CcpNMR (Vranken et al., 2005). For the relaxation dispersion curves, effective transverse relaxation rates *R*_2,eff_ (comprising intrinsic transverse relaxation rate *R*_2,0_ and any exchange contribution *R*_ex_) were extracted from experimental *R*_1ρ_ values using separately recorded *R*_1_ experiments (Palmer & Massi, 2006). For PRE experiments, ratios of peak intensities in spectra of para- and diamagnetic species (FhaC^220R1^ and FhaC^220R1dia^ for solid-state experiments, oxidized and reduced FhaC^195R1^ in case of the solution-state experiments, respectively) were calculated. These para- versus diamagnetic signal intensity ratios do not normalize to 1 in our case. In the solid-state experiments, this is most likely due to variations between the samples in terms of efficiency of protein reconstitution into liposomes and total amounts of sample transferred to the NMR rotor. Both in solid and solution state, spectroscopic factors likely also play a role (incomplete longitudinal relaxation and thus lower signal-to-noise in the spectra of diamagnetic samples due to the use of short inter-scan delays of 1 s (Iwahara et al., 2007)). We have thus opted to normalize PRE ratios to the maximum ratio observed in each experiment, which was always observed in one of the residues furthest from the paramagnetic center (Ile^136^ in FhaC^220R1^, Ile^14^ in FhaC^195R1^). This is equivalent to normalizing signals within each spectrum to a reference signal whose intensity is unaffected by PRE effects. We then only analyzed relative signal attenuation levels, instead of attempting to extract quantitative distance measures. Error bars of PRE intensity ratios were calculated based on spectral noise levels (root-mean-standard deviation of the spectral noise) using standard error propagation. To estimate expected distances between FhaC Ile residues and the MTSL label on residue 220, and consequently relative PRE attenuation levels, an ensemble of 200 MTSL conformations compatible with labeling on FhaC residue 220 was calculated using the mtsslSuite web server (Hagelueken et al., 2015; Hagelueken et al., 2012) (http://www.mtsslsuite.isb.ukbonn.de/) and the FhaC crystal structure (PDB 4QKY) with residue 220 changed to Cys in PyMOL (The PyMOL Molecular Graphics System. Schrödinger, LLC). The average position of the paramagnetic center (taken as halfway between nitrogen and oxygen atoms of the MTSL nitroxide ring) was calculated from the coordinates of these 200 conformations; distances from that position to Ile Cδ1 nuclei were calculated using PyMOL.

### EPR Experiments

PELDOR experiments were performed at Q-band frequency (∼34 GHz) using a Bruker EleXsys E580 spectrometer equipped with an overcoupled Bruker EN 5107D2 resonator. Pulses were generated with a Bruker SpinJet AWG and amplified with a 50 W TWT amplifier. The experiments were performed at 50 K and 30 K using a variable-temperature cryogen-free system (Oxford, Oxford, UK). The deadtime-free, four-pulse PELDOR sequence [(π/2)probe ― τ1 ― (π)probe ― τ1 + t ― (π)pump ― τ2 –t ― (π)probe ― τ2 ― (echo)] was employed with a 200-ns τ1 delay and τ2 delays ranging from 3,200 ns to 7,000 ns depending on the sample (Pannier et al., 2000). Probe pulses were 10 ns (π/2) and 20 ns (π) Gaussian-shaped pulses at a frequency corresponding to the maximum of the resonator response function and a magnetic field value corresponding to the high-field shoulder of the echo-detected field-swept spectrum. The pump pulse was implemented as a 24-ns pulse centered at a frequency 55 MHz higher than the probe frequency and corresponding to the maximum of the nitroxide field-swept spectrum. Raw time-domain PELDOR traces were background-corrected using the DeerAnalysis 2019 package (Jeschke et al., 2006), and the resulting signals were power-scaled in MATLAB to suppress sum and difference peaks arising from multispin effects. Distance distributions were then calculated from the scaled and background-corrected PELDOR traces by Tikhonov regularization. For FhaC^33R1+503R1^, FhaC^187R1+503R1^ and FhaC^195R1+503R1^, distance distributions were predicted using a pre-computed rotamer library of the MTSL spin probe attached to specific residues on the PDB structure (Jeschke, 2020).

### Mass fingerprinting of FhaC variants

Purified FhaC^C48+C224^ and FhaC^C195+C224^ variants were subjected to non-reducing SDS-PAGE, and acrylamide bands corresponding to the oxidized forms of the two proteins were excised. They were washed with 50 µL of acetonitrile (ACN)/NH_4_HCO_3_ (75/25) four times and dehydrated with ACN, or incubated in 10 mM DTT in NH_4_HCO_3_ for 30 min at 57°C and 30 min room temperature, followed by incubation in 55 mM iodoacetamide in 25 mM NH_4_HCO_3_ for 20 min in the dark, 3 washes with NH_4_HCO_3_ and dehydration with ACN performed twice. The pH of the samples was decreased to 2.0, digestion was performed with pepsin (0.01 µg/µL) (Promega, Charbonnieres-les- Bains, France) at a 1:50 enzyme:substrate ratio at 37°C for 3 hours, and the reaction was stopped by heating at 95°C for 10 min.

NanoLC-MS/MS analysis was performed using a nanoAcquity Ultra-Performance-LC (Waters, Manchester, UK) coupled to a Q-Exactive Plus Orbitrap mass spectrometer (Thermo Scientific, Illkirch, France). Peptides were trapped on a nanoACQUITY UPLC precolumn (C18, 180 µm x 20 mm, 5 µm particle size), and eluted from a nanoACQUITY UPLC column (C18, 75 µm x 250 mm, 1.7 µm particle size) at a constant temperature of 60°C. Mobile phases A and B were composed of 0.1% formic acid in water and 0.1% formic acid in ACN, respectively. Peptides were eluted with gradients of B from 1 to 8% for 2 min, 8 to 35% for 58 min, 35 to 90% for 1 min, 90% for 5 min, 90 to 1% B for 1 min and a concentration of 1% B for 20 min, with a constant flow rate of 400 nL/min. The source temperature of the mass spectrometer was set to 250°C and the spray voltage at 1.8 kV. Full scan MS spectra were acquired in positive mode with a resolution of 140,000, a maximum injection time of 50 ms, and an AGC target value of 3×10^6^ charges. The 10 most intense multiply charged peptides per full scan were isolated using a 2 m/z window and fragmented using higher energy collisional dissociation (normalized collision energy of 27). MS/MS spectra were acquired with a resolution of 17,500, a maximum injection time of 100 ms and an AGC target value of 1 x 10^5^, and dynamic exclusion was set to 60 sec. The system was fully controlled by XCalibur software v3.0.63, 2013 (Thermo Scientific) and NanoAcquity UPLC console v1.51.3347 (Waters). The MS/MS data were interpreted using a local Mascot server with MASCOT 2.5.0 algorithm (Matrix Science, London, UK). Spectra were searched with a mass tolerance of 5 ppm for MS and 0.07 Da for MS/MS data, using none as enzyme. Oxidation (+15.99 Da), and carbamidomethylation (57.02 Da) were specified as variable modifications. Protein identifications were validated with a Mascot ion score above 25.

### Native MS and ion mobility

Purified FhaC was buffer exchanged into 100 mM ammonium acetate buffer, pH 6.8, supplemented with 50 mM bOG using a P6 desalting column (Biorad, Marnes-la-Coquette, France). Samples were directly infused with nano-electrospray ionization with in-house-prepared gold-coated borosilicate glass capillaries with a spray voltage of +1.4 kV. Spectra were recorded on a quadrupole TOF instrument (Synapt G2 HDMS with 32K quadrupole, Waters) optimized for transmission of native, high-m/z protein assemblies. Critical voltages and pressures throughout the instrument were 50 V, 10 V, 150 V and 15 V for the sampling cone, extraction cone, trap and transfer collision cell, respectively, with pressures of 9 mbar, 1.47 × 10^−2^ mbar and 1.21 × 10^−2^ mbar for the source, trap and transfer regions unless indicated otherwise. CIU ion mobility experiments were performed with 50 V sampling cone; 50-200 V trap collision energy; 42 V trap DC bias; and 15 V transfer collision energy. Pressures throughout the instrument were 9 and 1.46 × 10^−2^ mbar for the source and trap/transfer collision cells. All spectra were processed with Masslynx v4.1 (Waters). Collision cross section calibration was performed using GDH, ADH, ConA and PK as proteins standard as described (Allen et al., 2016). It should be noted that due to the generally lower charge states observed for membrane proteins, and the increased collision energies required (compared to soluble proteins) for gentle release of proteins from detergent micelles, the CCS values reported here are less accurate and intended for qualitative comparison rather than quantitative matching to theoretical models.

### Peptide binding assays

Synthesized peptides were dissolved in DMSO to a final concentration of 100 mM and added to the protein sample at final concentrations of 10 μM FhaC and 100 μM peptide. To correct for non-specific and detergent-specific binding, SphB1-αβ was run at identical concentrations and conditions. For both proteins the fraction of peptide-bound protein was calculated based on peak intensities, after which the binding to the decoy protein was subtracted to correct for non-specific binding.

## Acknowledgements

We thank Xavier Hanoulle for useful suggestions and discussions at the start of this project and Isabelle Landrieu for her support and advice. This work was funded by the Agence Nationale de Recherche grant ANR-17-CE11-0043-02 “OPEN_BAR” to FJD and a “Projets exploratoires premier soutien (PEPS)” grant by the CNRS and the University of Lille to RS. Financial support and spectrometer access by the IR-RMN-THC FR 3050 CNRS for conducting NMR experiments at the IR-RMN platforms in Lille and Grenoble is gratefully acknowledged. EPR experiments were performed within the national facility RENARD at the University of Lille (Federation IR-RPE 3443). Support by the Centre National de la Recherche Scientifique (CNRS), Université de Strasbourg (Unistra) and the French Proteomic Infrastructure (ProFI; ANR-10-INBS-08-03) is acknowledged. The authors would also like to thank the IdeX program of the University of Strasbourg for funding the Synapt G2Si instrument.

---

**Figure 3 Supplement 1.**
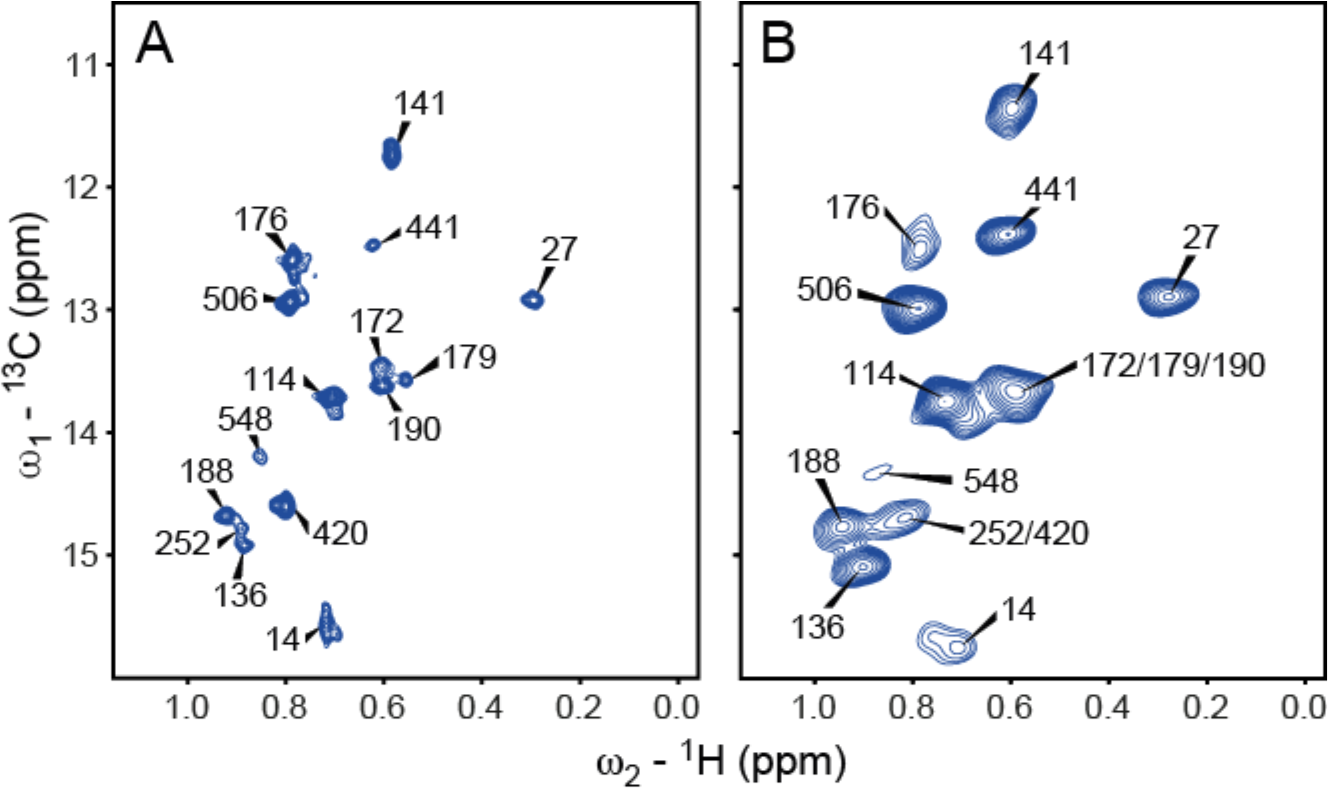
NMR analyses of Ile δ_1_ methyl labeled FhaC in lipid bilayers. (A) Methyl region of a solution-state heteronuclear multiple quantum coherence (HMQC) ^13^C-^1^H correlation spectrum of u-(^2^H, ^15^N), Ile-δ1(^13^CH3)-labeled FhaC^195R1^ in ^2^H-MSP1D1 nanodiscs prepared from deuterated (d_54_-) DMPC and DMPG (2:1) lipids, recorded on a 900 MHz spectrometer. The MTSL tag on residue 195 was reduced with ascorbic acid; peak positions are identical to those of wt FhaC in nanodiscs. (B) Same region of a scalar coupling-based solid-state J-HSQC ^13^C-^1^H correlation spectrum of wt FhaC (same isotope labeling as in (A)) in d_54_-DMPC/DMPG liposomes, recorded on an 800 MHz spectrometer at 50 kHz MAS.

**Figure 3 Supplement 2.**
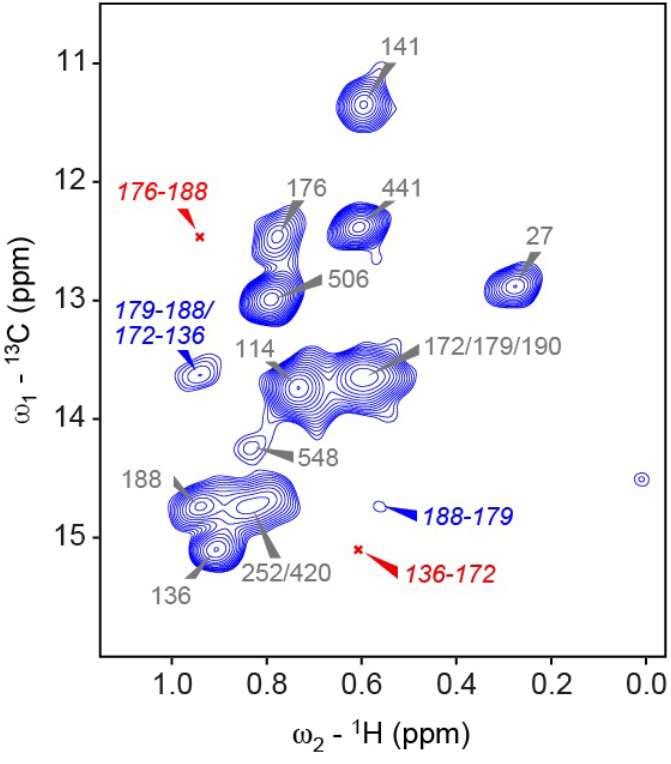
NMR analysis of through-space contacts between Ile δ_1_ methyl groups in FhaC in liposomes. 2D hChH correlation spectrum with 6.4 ms RFDR (Bennett et al., 1992) ^1^H-^1^H mixing of FhaC u-(^2^H, ^15^N), Ile-δ_1_(^13^CH_3_) in deuterated (d_54_-) DMPC/ DMPG liposomes, recorded on an 800 MHz NMR spectrometer at 50 kHz MAS, to visualize through-space correlations between Ile δ_1_ methyl groups close in space. Among expected inter-residue cross-peaks (^1^H-^1^H distance below 6 Å), peaks present in the spectrum are indicated in blue, those which are absent in red.

**Figure 3 Supplement 3.**
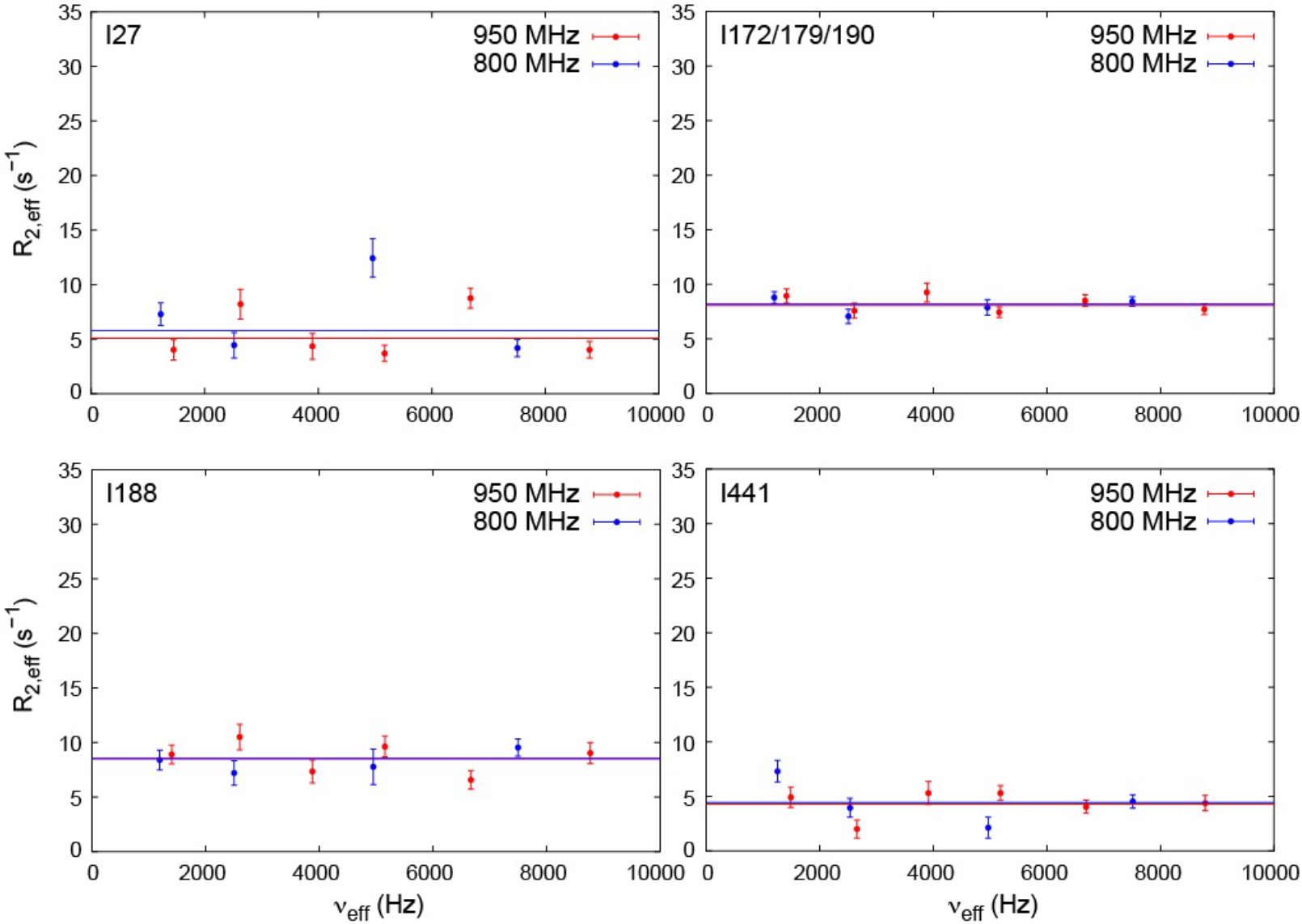
NMR relaxation dispersion experiments to measure µs time scale exchange dynamics in FhaC. Effective ^13^C transverse relaxation rates *R*_2,eff_ extracted from solid-state NMR *R*_1ρ_ relaxation dispersion experiments (Lewandowski et al., 2011; Ma et al., 2014) on selected Ile-δ_1_ methyl groups of u-(^2^H, ^15^N), Ile-δ_1_(^13^CH_3_)-labeled wt FhaC, recorded on 800 (blue) and 950 MHz (red) spectrometers at 50 kHz MAS frequency and 17°C sample temperature. Horizontal lines are best fits to the data using a model of no exchange (i.e. constant *R*_2,eff_ values for varying applied *B*_1_ radiofrequency fields and thus varying effective fields ν_eff_). Models assuming exchange do not fit the data significantly better according to F test statistics or Akaike’s information criterion (AIC) in any of the Ile- δ_1_(^13^CH_3_) groups of FhaC. Notably, data from residue Ile^548^ in strand β16, at the barrel junction with strand β1, could not be reliably analyzed due to low signal-to-noise.

**Figure 3 Supplement 4.**
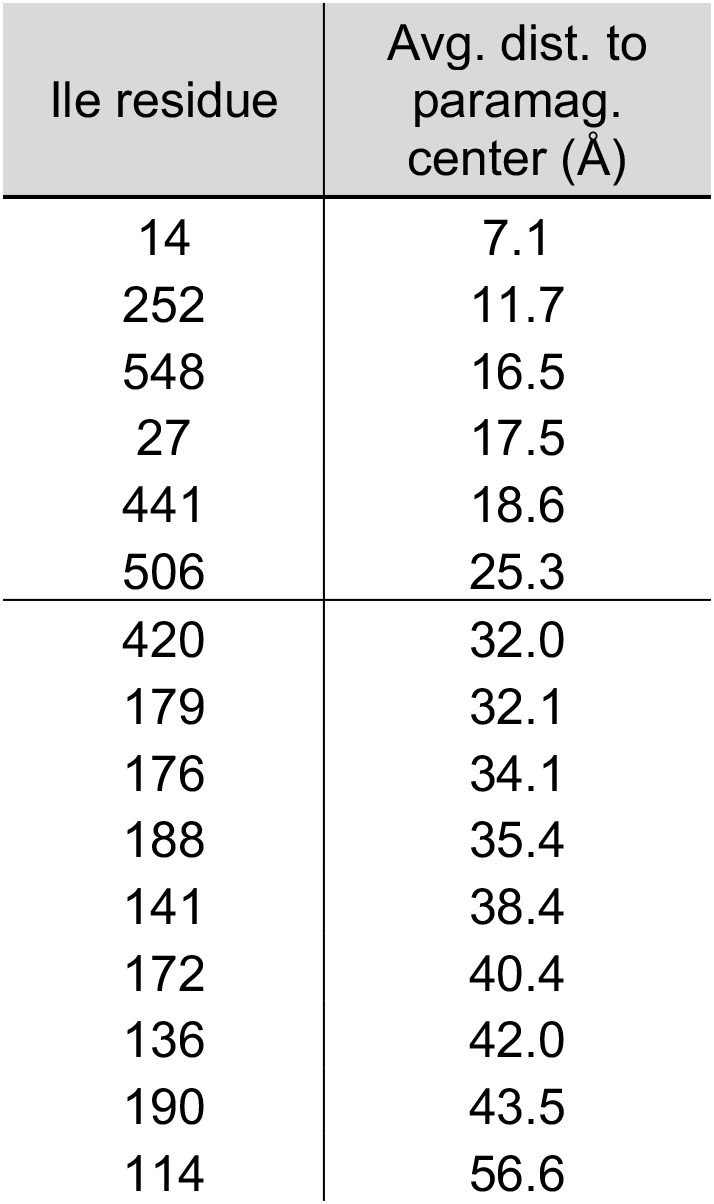
Estimated Ile Cδ_1_ – MTSL distances in the crystal structure conformation of FhaC^220R1^. Shown are distances (in Å) between Ile Cδ_1_ nuclei and the estimated average position of the paramagnetic center in FhaC with a MTSL spin label on residue 220 (FhaC^220R1^). An ensemble of 200 MTSL conformations compatible with labeling on FhaC residue 220 was calculated using the mtsslSuite web server (Hagelueken et al., 2012; Hagelueken et al., 2015) (http://www.mtsslsuite.isb.ukbonn.de/) and the FhaC crystal structure (PDB 4QKY). The average position of the paramagnetic center (taken as halfway between nitrogen and oxygen atoms of the MTSL nitroxide ring) was calculated from the coordinates of these 200 conformations; distances from that position to Ile Cδ_1_ nuclei were calculated using PyMOL (The PyMOL Molecular Graphics System. Schrödinger, LLC). A horizontal line in the table indicates the distance from the paramagnetic center up to which attenuation effects on the NMR resonance of the corresponding Ile residue are expected if FhaC assumes a conformation as in the crystal structure.

**Figure 4 Supplement 1.**
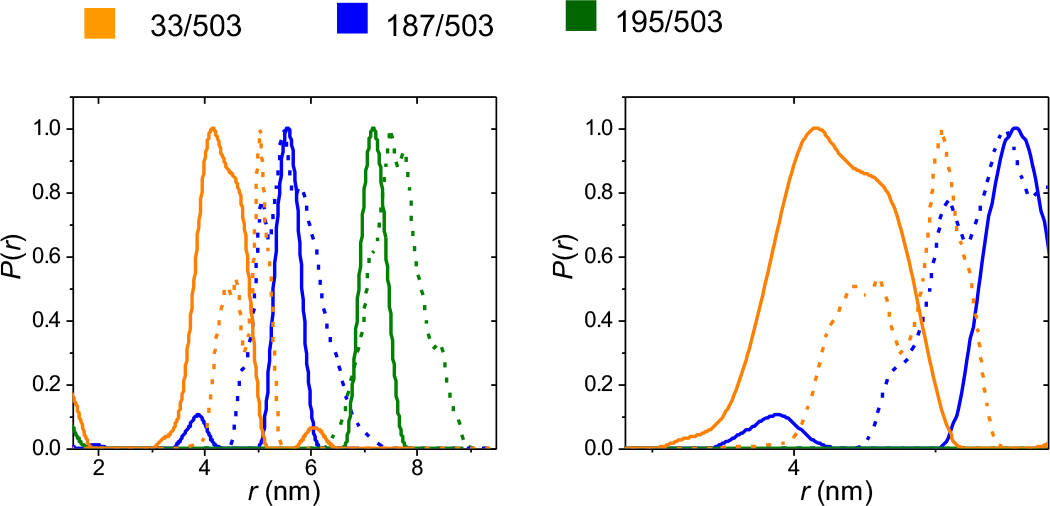
Distance distributions from PELDOR experiments. The distance distributions obtained for FhaC^33R1+503R1^ (orange), FhaC^187R1+503R1^ (blue) and FhaC^195R1+503R1^ (green) in bOG (solid lines) are compared with those predicted using a pre-computed rotamer library of the MTSL spin probe attached to specific residues on the PDB structure of FhaC (dashed lines) (Jeschke, 2020). In the right panel, a zoom on the 3-5 nm region shows the broad distribution for FhaC^33R1+503R1^.

**Figure 4 Supplement 2.**
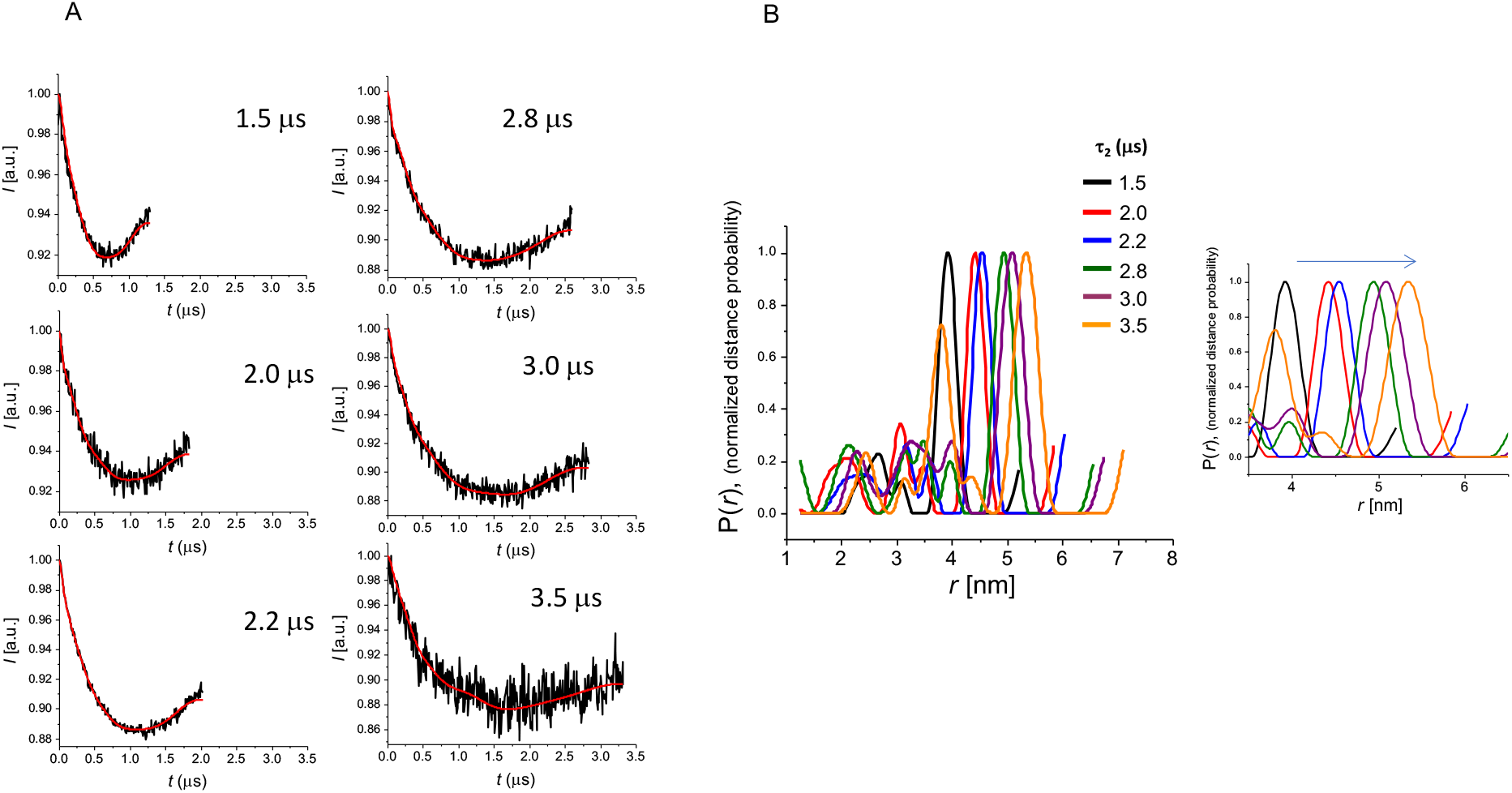
Dipolar evolution signals recorded with different delays τ*_2_* and corresponding distance distributions. FhaC^195R1+503R1^ in *E. coli* lipids liposomes was used in this experiment. (A) The dipolar evolution signals were measured at increasing dipolar evolution times *t*. (B) The longest distance measured shifts to longer values for longer dipolar evolution times *t* since long, but not short distances are sensitive to the value used in PELDOR experiments. The lipid environment decreases the dipolar evolution time that can be applied, which results in an apparent shift to smaller distance distribution values. The right panel is a zoom on the 4-6 nm region.

**Figure 4 Supplement 3.**
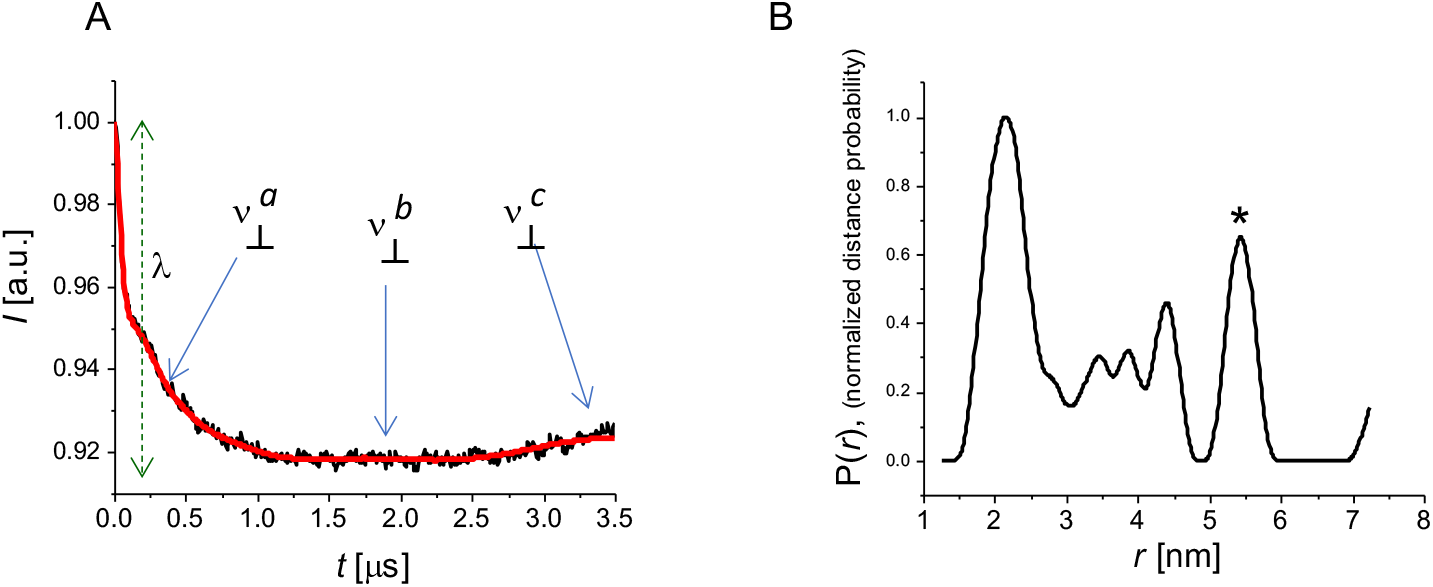
Effect of the L6-barrel interaction on conformational changes of FhaC. (A) Dipolar evolution function for the PELDOR signal for FhaC^R492+195R1+503R1^. (B) Distance distribution obtained by Tikhonov regularization of the signal depicted in (A). The asterisk corresponds to the longest distance measurable as a function of the *t* parameter applicable in this experiment.

**Figure 4 Supplement 4.**
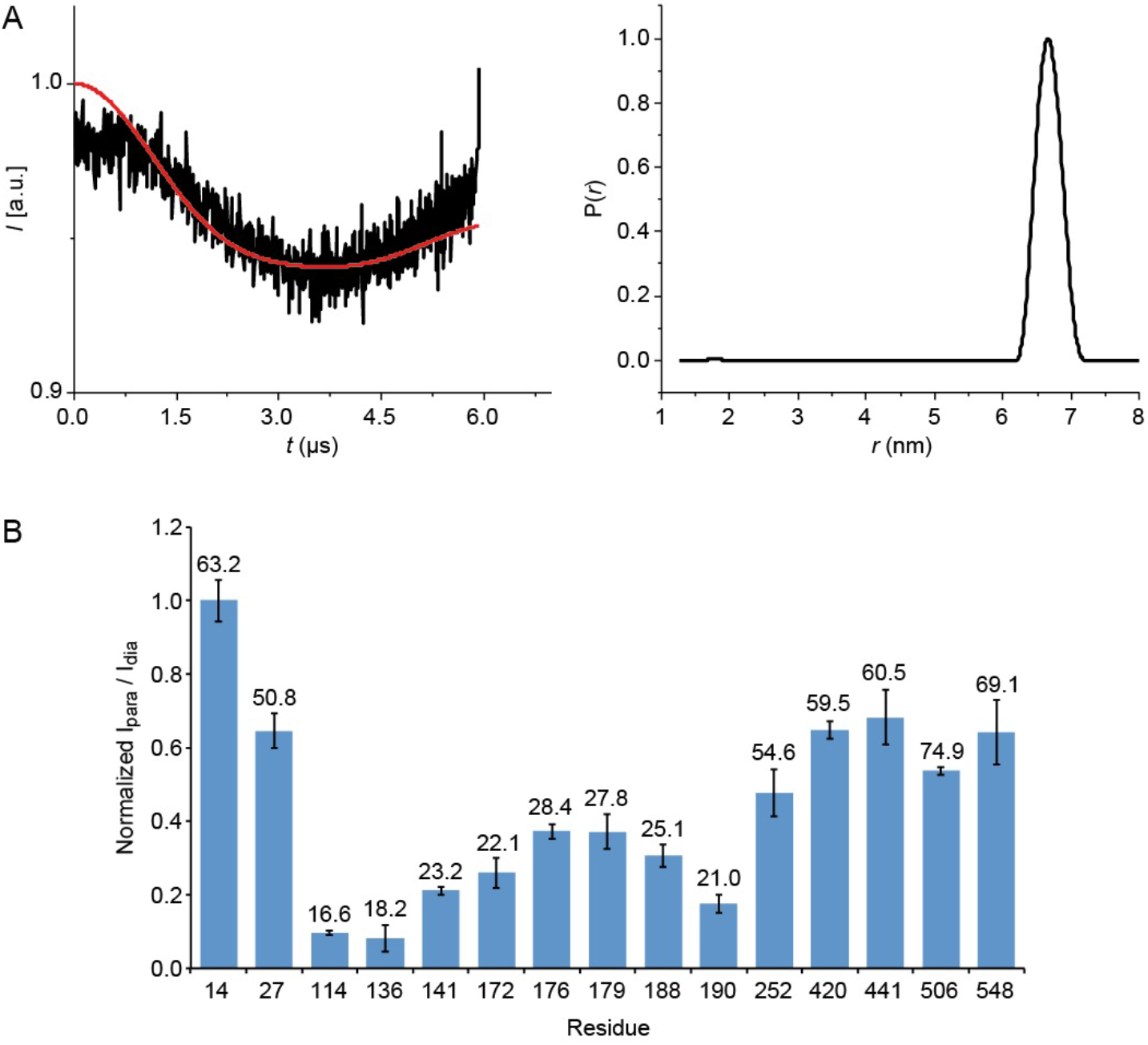
Spectroscopic analyses of FhaC in nanodiscs. (A) Dipolar evolution function (*left*) and Tikhonov regularization (*right*) of the PELDOR signal of FhaC^195R1+503R1^ in nanodiscs. (B) NMR paramagnetic relaxation enhancement experiments on FhaC^195R1^ in nanodiscs. Ratios *I*_para_/*I*_dia_ of paramagnetic vs. diamagnetic FhaC^195R1^ Ile-δ_1_ methyl peak intensities (normalized to their maximum value found for Ile^14^; see Methods) extracted from ^13^C-^1^H heteronuclear multiple-quantum coherence (HMQC) experiments in solution before and after reduction of the MTSL spin label by addition of a 10-fold molar excess of ascorbic acid. Spectra were recorded on a 900 MHz spectrometer at 32°C. Error bars are calculated based on spectral noise levels. Numbers above the bars indicate the distance between the Cδ_1_ nucleus of the corresponding Ile residue and the average position of the paramagnetic center in an ensemble of conformations of the MTSL tag attached to Cys^195^ in the FhaC crystal structure, calculated using the MtsslWizard PyMOL plugin (Hagelueken et al., 2012). Relative levels of signal attenuation due to the MTSL tag are perfectly in line with the relative distances of the corresponding residues from the paramagnetic center modeled onto the FhaC crystal structure, suggesting that FhaC in nanodiscs does not populate alternative conformations.

**Figure 5 Supplement 1.**
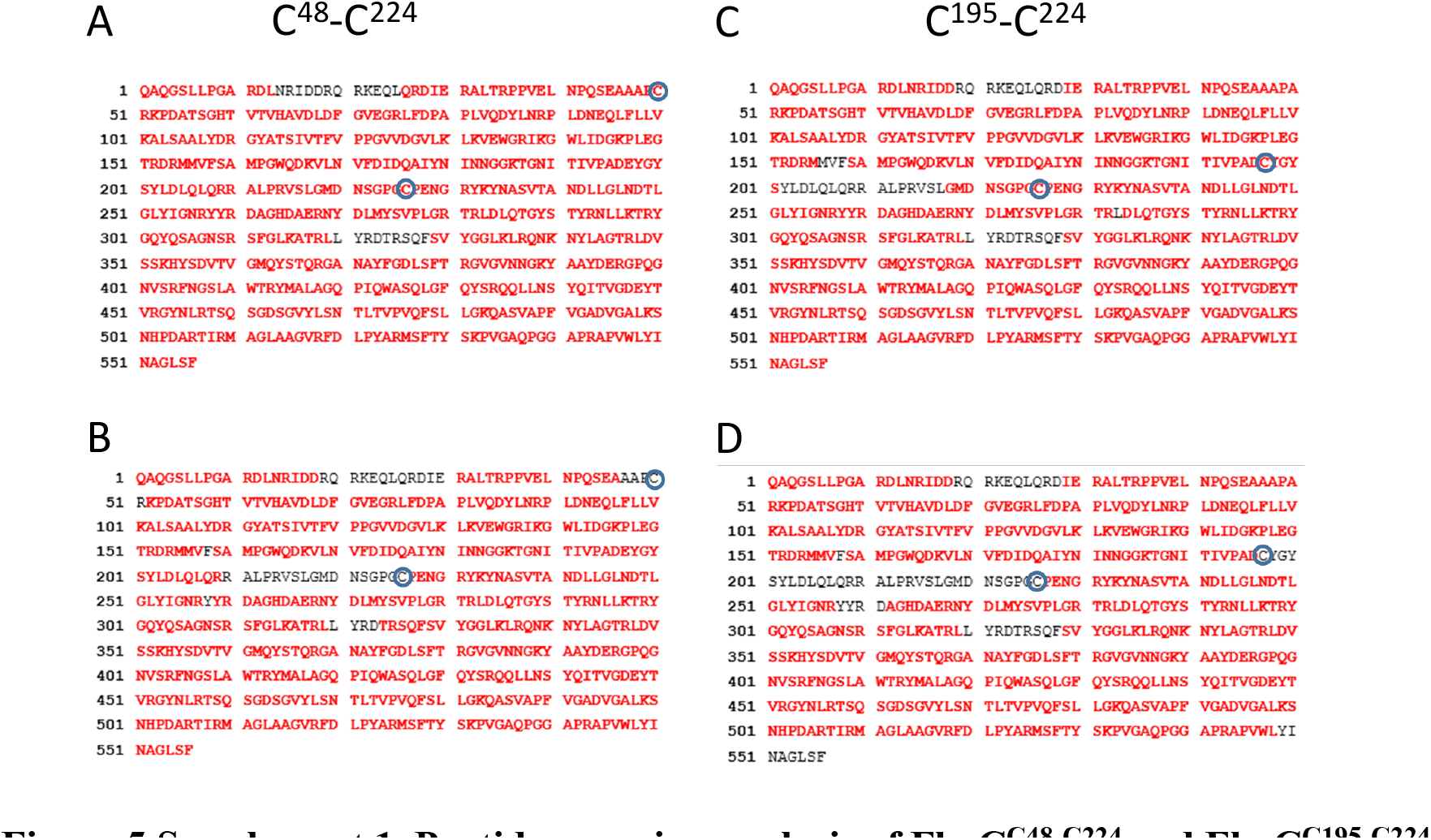
Peptide mapping analysis of FhaC^C48-C224^ and FhaC^C195-C224^. Residues in red represent sequence coverage with (A,C) or without (B,D) reduction and alkylation. In the latter cases, the regions that contain the Cys residues were not characterized, suggesting the presence of an intramolecular S-S bond in both variants. Note that the sequences shown here contain an N-proximal Gly-Ser insertion for cloning purposes that has no effect on the structure or the activity of FhaC. The numbering of FhaC throughout the text corresponds to that of the native protein without this insertion.

**Figure 6 Supplement 1.**
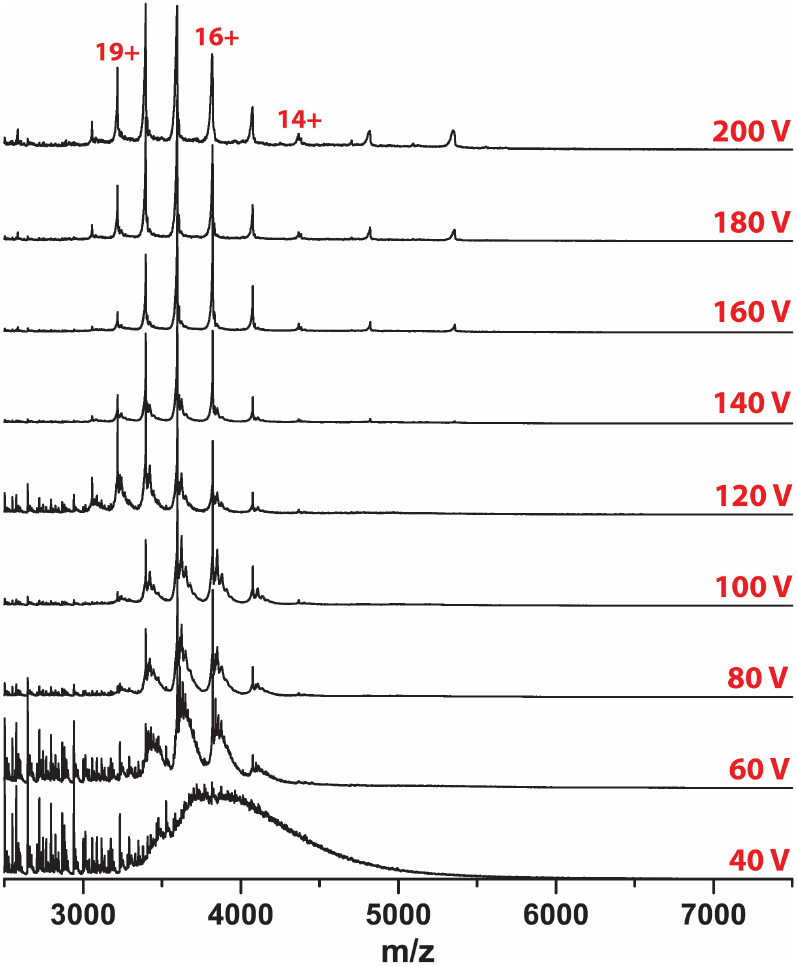
Native MS analysis of FhaC in bOG micelles. The spectra were obtained at increasing collisional energy.

**Figure 6 Supplement 2.**
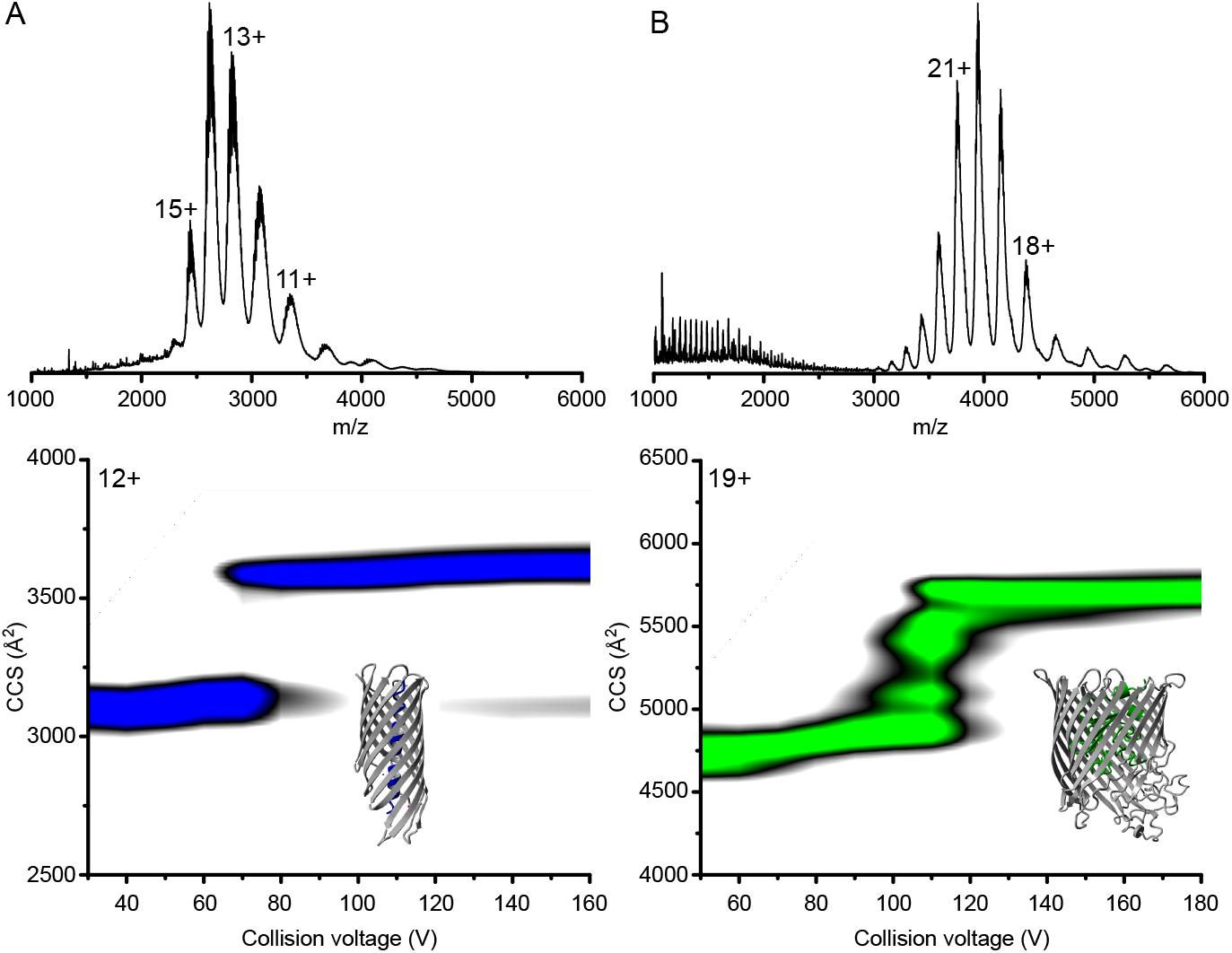
Mass spectra and CIU plots of control OMPs. (A) SphB1-*αβ*is a truncated autotransporter (AT) containing only the *β* barrel with the preceding helical linker inserted in the barrel pore. (B) The TonB-dependent transporter BfrG is composed of a *β* barrel with a soluble N-terminal plug domain inserted in the barrel. The structural models shown are those of related transporters (PDB 1UYN and 3QLB, respectively), as the structures of SphB1-*αβ* and BfrG are not available. The mass spectra of the two OMPs released from their bOG micelles are shown at the top, and the CIU plots are below. Both show a single CIU transition, which suggests that the *β* barrels remain intact, while the soluble domains are ejected and unfold.

**Figure 6 Supplement 3.**
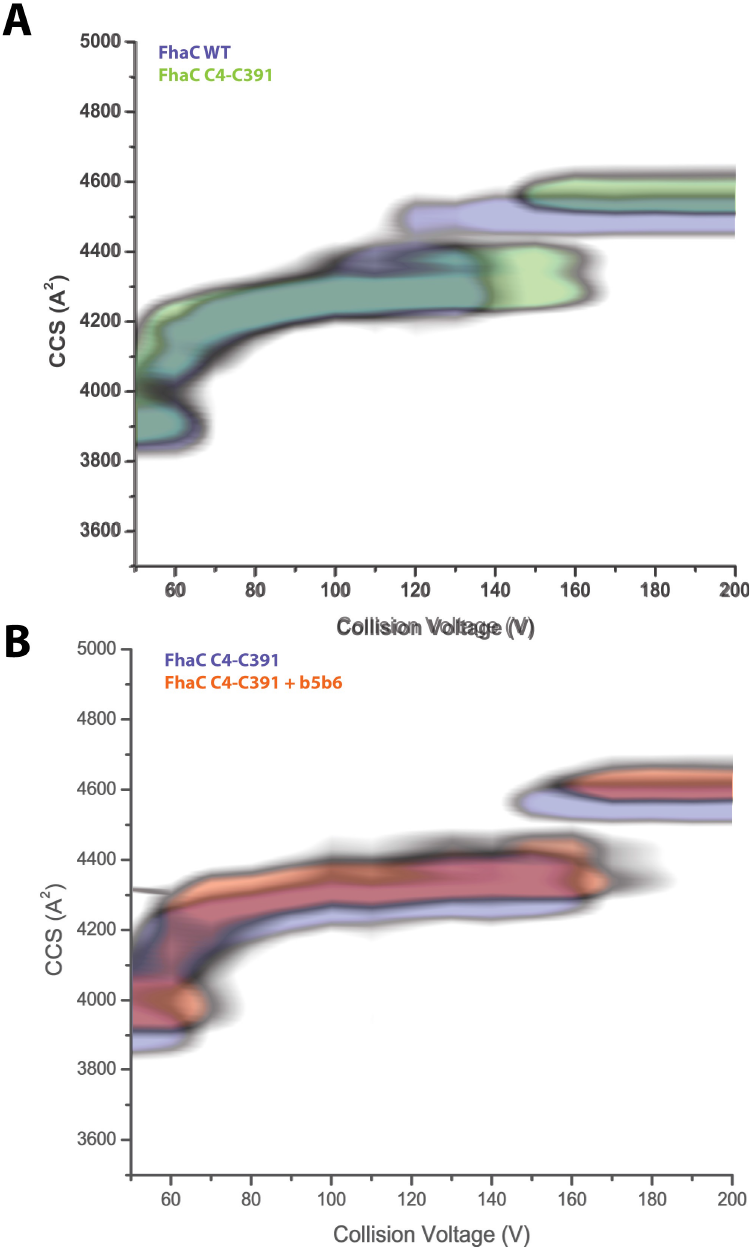
CIU plots of FhaC^C4+C391^. (A) Comparison of the CIU plots of wt FhaC (blue) and the FhaC^C4+C391^ variant (green). (B) Overlay of the CIU plots of unbound FhaC^C4+C391^ (blue) and FhaC^C4+C391^ with the b5-b6 peptide bound (red). As for wt FhaC (see Figure 7), binding of the peptide to FhaC^C4+C391^ increased CCS values at both low and high CE, suggesting that it induces enlargement of the *β* barrel, even with H1 inside.

**Figure 7 Supplement 1.**
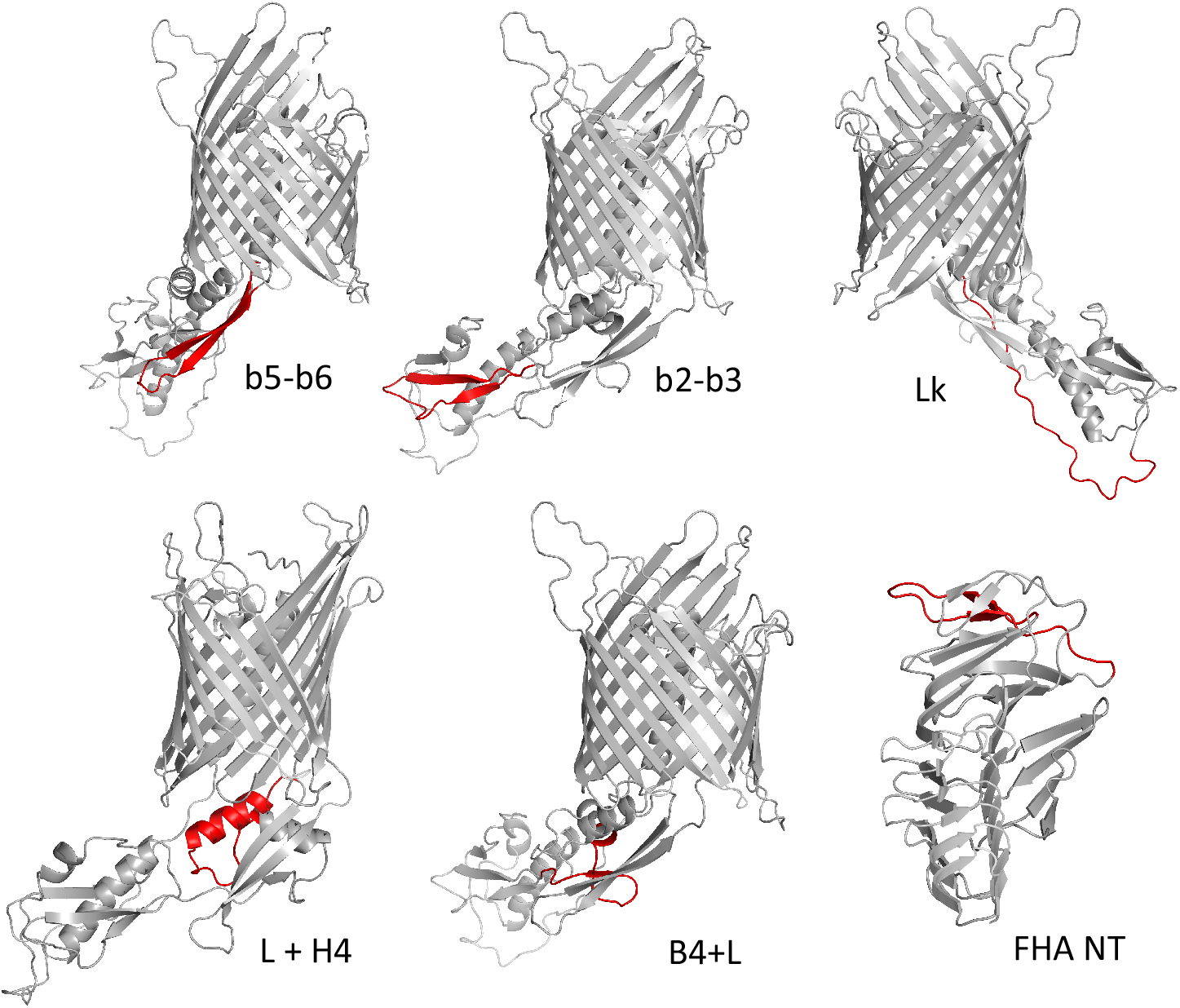
Synthetic peptides used in this study. The first 5 peptides are shown in red on the structural model of FhaC, and the last one on the structural model of the N-terminal portion of FhaB (PDB 1RWR). The b5-b6 peptide (GKTGNITIVPADEYGYSYLDLQLQR) corresponds to the last two β strands of the POTRA2 domain that form an amphipathic β hairpin immediately preceding Β1, the first strand of the β barrel. The b2-b3 peptide (SIVTFVPPGVVDGVLKLKVEWGR) encompasses the last two β strands of the POTRA1 domain. The Lk peptide (RPPVELNPQSEAAAPARKPDATSGH) corresponds to the linker between the H1 helix and the POTRA1 domain. The L+H4 peptide (AMPGWQDKVLNVFDIDQAIYNINNG) encompasses the loop (extended) region that precedes the H4 α helix and the H4 helix of the POTRA2 domain. The B4+L peptide (RIKGWLIDGKPLEGTRDR) corresponds to the β strand b4 of the POTRA2 domain followed by a loop region. Finally, the FHA-NT peptide (QTQVLQGGNKVPVVNIADPNS) corresponds to the N-terminal β strands b2 and b3 of FhaB forming a short hairpin, preceded and followed by loop regions.

**Figure 7 Supplement 2.**
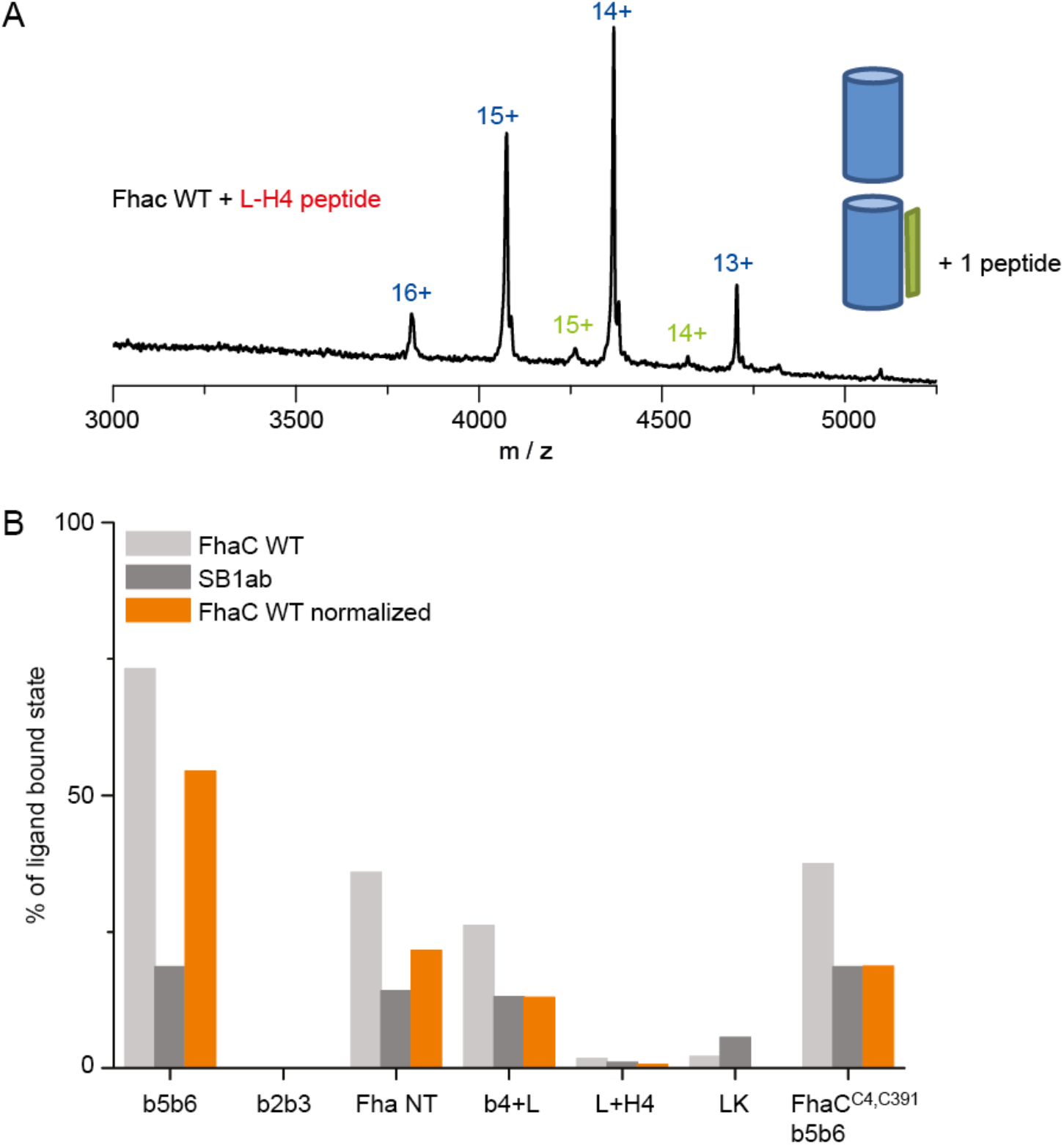
Binding of synthetic peptides to FhaC. (A) Under similar conditions as in Fig. 7A (collisional energy of 150 V), only minimal binding was detected in the mass spectrum of FhaC incubated with the L+H4 peptide. (B) Quantification of the binding of synthetic peptides to FhaC (light grey) and to the control β-barrel protein SphB1-αβ (SB1ab, dark grey; used to correct for non-specific binding). Orange bars show normalized values.

**Figure 7 Supplement 3.**
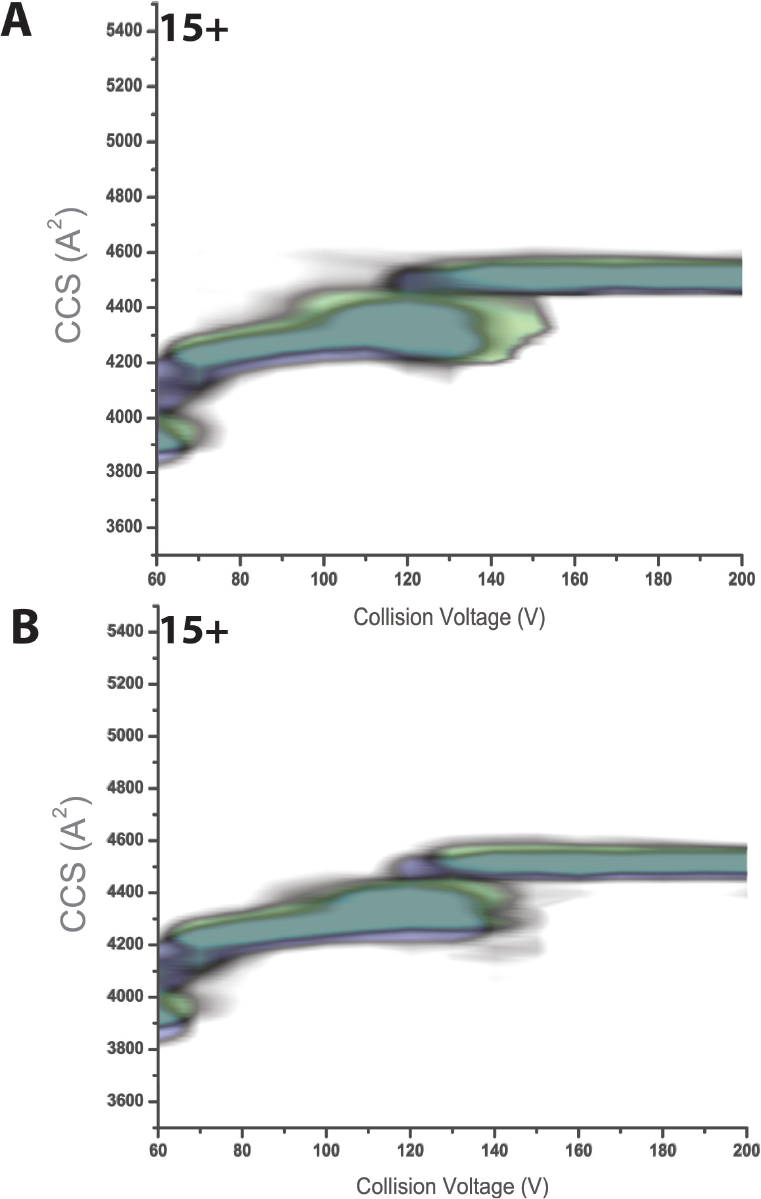
CIU plots of FhaC incubated with synthetic peptides. (A) FhaC without (blue) or with (green) the Fha-NT peptide. (B) FhaC without (blue) or with (green) the B4+L peptide. In both cases the presence of the peptide caused an increased CCS at low collision voltage, but not at elevated collisional activation.

**Figure 7 Supplement 4.**
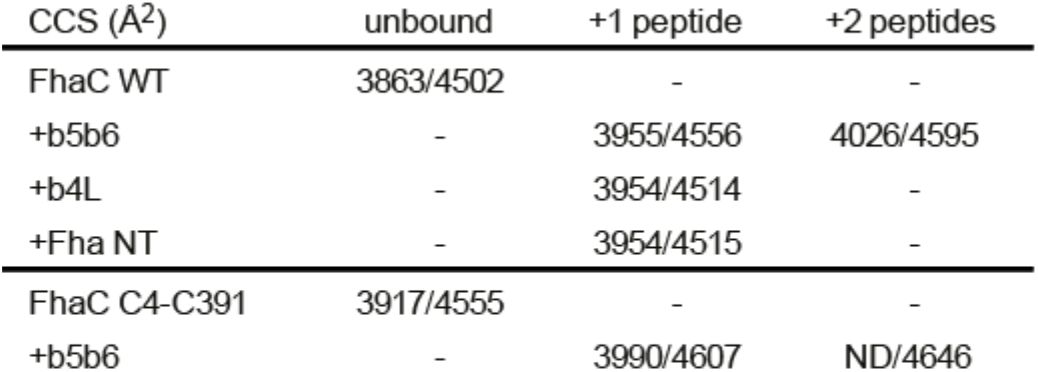
CCS of FhaC with various peptides. determined at low and high CE (listed before and after the slash). The measured values indicate that only b5-b6 enlarges FhaC in both conditions.

## Notes

### Competing Interest Statement

The authors have declared no competing interest.

